# Targeting SUV4-20H epigenetic enzymes as therapeutic strategy for enhancing topoisomerase II poisoning in prostate cancer

**DOI:** 10.1101/2025.03.11.642623

**Authors:** Fatima Alhourani, Marine Tauziet, Julie Patouillard, Benoit Miotto, Véronique Baldin, Aurélie Gennetier, Simon George, Xavier Miahle, Cyril Ribeyre, Philippe Pourquier, Florence M. Cammas, Adeline Torro, Diego Tosi, Céline Gongora, Mona Dergham, Raghida Abou-Merhi, Eric Julien

## Abstract

DNA topoisomerase II (TOP2) plays a crucial role in DNA-associated processes by inducing transient DNA double-strand breaks, making it an important target for DNA-damage stabilizing agents. Commonly used in cancer therapy, these agents are designed to interfere with TOP2 cleavage complexes on chromatin. However, the epigenetic pathways influencing their effectiveness and the resultant cellular responses remain elusive. Here, we combine *in vitro* as well as *in vivo* genetic and pharmacological approaches in prostate cancer to demonstrate that inhibiting the histone H4-lysine 20 (H4K20) methyltransferases SUV4-20H1 and SUV4-20H2 induces synthetic lethality when combined with TOP2 poisons, such as etoposide. Remarkably, we show that the loss of SUV4-20H enzymes, which prevents the conversion of H4K20 mono-methylation to higher methylation states, has minimal impact on prostate cancer cell behavior under normal conditions. However, these innocuous epigenetic changes significantly enhances the trapping of TOP2 complex in chromatin and increases DNA damage in response to etoposide. Furthermore, SUV4-20H depletion impairs the repair of TOP2-induced DNA breaks by disrupting the switch from RPA to RAD51 foci at damage sites, leading to extensive cancer cell death and inhibition of prostate tumor growth. Overall, these findings suggest that dual targeting of SUV4-20H and TOP2 activity on chromatin represents a promising therapeutic strategy for prostate cancer, where SUV4-20H2 emerges as a potential marker of aggressive disease and high metastatic risk.

## INTRODUCTION

The human type-II topoisomerases (TOP2) are nuclear enzymes that are essential for the removal DNA topological constraints that arise during DNA replication, transcription and mitosis ^1,2^. TOP2 acts as a transient cleavage complex intermediates (TOP2ccs) capable of generating reversible double strand DNA breaks (DSBs) by cleaving the DNA phosphodiester backbone ^1^. This TOP2ccs complex consists of covalent phosphotyrosyl bonds of TOP2 monomers at the 5′ end of the breaks in both strands. After passing another molecule of DNA through the break site, TOP2 re-ligates the ends and subsequently dissociates from DNA ^3^. While TOP2ccs are normally reversible and short-lived under physiological conditions, they can be stabilized and trapped by specific poisons, such as etoposide or mitoxantrone ^3,4^. These poisons inhibit the re-ligation step of the reaction, leading to irreversible and cytotoxic TOP2-linked DNA breaks ^5^. Therefore, TOP2 poisons are commonly used in oncology to treat a large range of malignancies, such as lung, breast, testicular and castration-resistant prostate cancer ^4,6,7^. However, despite their relative effectiveness, the use of TOP2 poisons is often associated with important side effects, including long-term toxicities, secondary malignancies, and dose-limiting cardiotoxicity ^6,8^. Thus, identifying alternative strategies aimed at mitigating the dosage of TOP2 poisons while increasing their effectiveness are important issues from both fundamental and clinical perspectives.

The methylation of histone H4 at lysine 20 (H4K20) is an abundant cell-cycle regulated epigenetic modification that can exist as three distinct states: mono- (H4K20me1), di- (H4K20me2) or tri-methylated (H4K20me3) ^9,10^. The lysine methyltransferase SET8 is the sole enzyme responsible for H4K20me1, which can be subsequently converted to H4K20me2 and H4K20me3 by the lysine methyltransferases SUV4-20H1 and SUV4-20H2, collectively referred as SUV4-20Hs^9^. Through their ability to induce H4K20me2 and H4K20me3 signals, SUV4-20Hs enzymes have been associated with the regulation of heterochromatin, DNA replication and repair mechanisms ^11–15^. Yet, SUV4-20H2 knockout mice exhibit no apparent phenotype, suggesting non-essential functions of this enzyme during development and in normal adult tissues ^16^. In contrast, SUV4-20H1 knockout leads to developmental defects and perinatal lethality likely caused by multiple chromatin alterations ^16^.

While our understanding of the physiological functions of SUV4-20H enzymes is still in its infancy, several reports suggested a pathological role of these enzymes in several cancers ^17^. While its depletion can drive malignant transformation in *TP53*-deficient adult stem cells, SUV4-20H1 expression could inhibit tumor growth in a mouse glioblastoma xenograft model ^18,19^. However, SUV4-20H1 is also amplified and overexpressed in other cancer types ^17^, reflecting a more complex pathological role than previously thought. Similarly, reduced SUV4-20H2 expression is observed in breast cancer and correlates with EGFR resistance in lung cancers ^20–22^, while its overexpression has been shown to promote hepatocarcinoma progression ^23^. Interestingly, however, targeting both SUV4-20H1 and SUV4-20H2 sensitize liver cancer cells to Poly(ADP-ribose) polymerase (PARP) inhibition, suggesting that these enzyme might constitute attractive targets for new anti-cancer strategies ^23^.

Herein, we identify SUV4-20H2 as a marker of aggressiveness in prostate cancer independently of the expression of SUV4-20H1 and that is not essential for the survival and proliferation of prostate cancer cells. However, inhibiting both SUV4-20H enzymes and altering H4K20me levels across the genome significantly increases the chromatin trapping of TOP2ccs in response to etoposide. Additionally, the loss of SUV4-20H enzymatic activity disrupts the repair of TOP2ccs-induced DNA breaks by preventing the focal accumulation of RAD51 at damage sites, leading to extensive cancer cell death and halting tumor growth in vivo. Thus, targeting SUV4-20Hs chromatin activity presents a powerful and novel epigenetic strategy to enhance the efficacy of TOP2 poisons in cancer therapy.

## RESULTS

### *SUV4-20H2* gene expression is a marker of aggressive prostate cancer

Immunohistology experiments have highlighted some alterations in the levels of histone H4K20 methylation in prostate tumors ^24^, suggesting an involvement of histone H4K20 methyltransferases in the etiology or the progression of this disease. To delve deeper into this hypothesis, we first examined the expression levels of the three main histone H4K20 methyltranferases—SET8, SUV4-20H1 and SUV4-20H2— utilizing the transcriptomic data set of the well-established prostate tumors cohort of the Cancer Genome Atlas (TCGA). This cohort encompasses 497 primary tumor samples and 52 normal samples serving as non-tumoral controls ^25^. No significant differences were observed in the expression levels of *SET8* and *SUV4-20H1* encoding genes (Figure 1A). In contrast, primary prostate tumors exhibited significantly elevated *SUV4-20H2* gene expression compared to non-tumoral samples (Figure 1A). However, in contrast to previously thoughts^26^, the up-regulation of *SUV4-20H2* in primary prostate tumors was independent of the Gleason score that currently define tumor grades (Supplementary Figure S1A). It was slightly more pronounced in younger patients (Supplementary Figure S1B) and associated with specific genetic features (Supplementary Figures S1C to S1G). While no significant correlation was identified with TMPRSS2 structural variant and SPOP mutation (Supplementary Figure S1C and S1D), heightened *SUV4-20H2* expression tended to coincide with P53 mutation (Supplementary Figure S1E), neoplasm disease lymph node stage 1 (Supplementary Figure S1F) and the absence of structural variants of the ERG oncogene (Supplementary Figure S1G). The expression of *SUV4-20H2* gene also correlated with genome instability, as indicated by the higher hypoxia score and the significant increase in the aneuploidy and the MSI mantis score in primary tumors samples with heightened *SUV4-20H2* expression (Supplementary Figure S2). In agreement with these results, *SUV4-20H2* mRNA levels were found associated with shorter progression free and disease-free survival (Figure 1B), suggesting that *SUV4-20H2* expression is a potential marker of aggressiveness in prostate tumors. Supporting this hypothesis, the increase in *SUV4-20H2* expression was linked to the occurrence of metastasis in two other distinct cohorts of PCa patients (Figure 1C). Furthermore, compared to all types of castration-resistant prostate cancer, high *SUV4-20H2* expression emerged as a marker of neuroendocrine prostate cancer (NEPC) (Figure 1D), which represents a highly aggressive subtype of prostate cancer that commonly emerges through a trans-differentiation process and evades conventional therapies. Altogether, these clinical data indicate that the up-regulation of *SUV4-20H2* expression is a new marker for prostate tumors associated with specific genetic alterations and that are prone to evolve towards metastatic and resistance stages.

**Figure 1:**
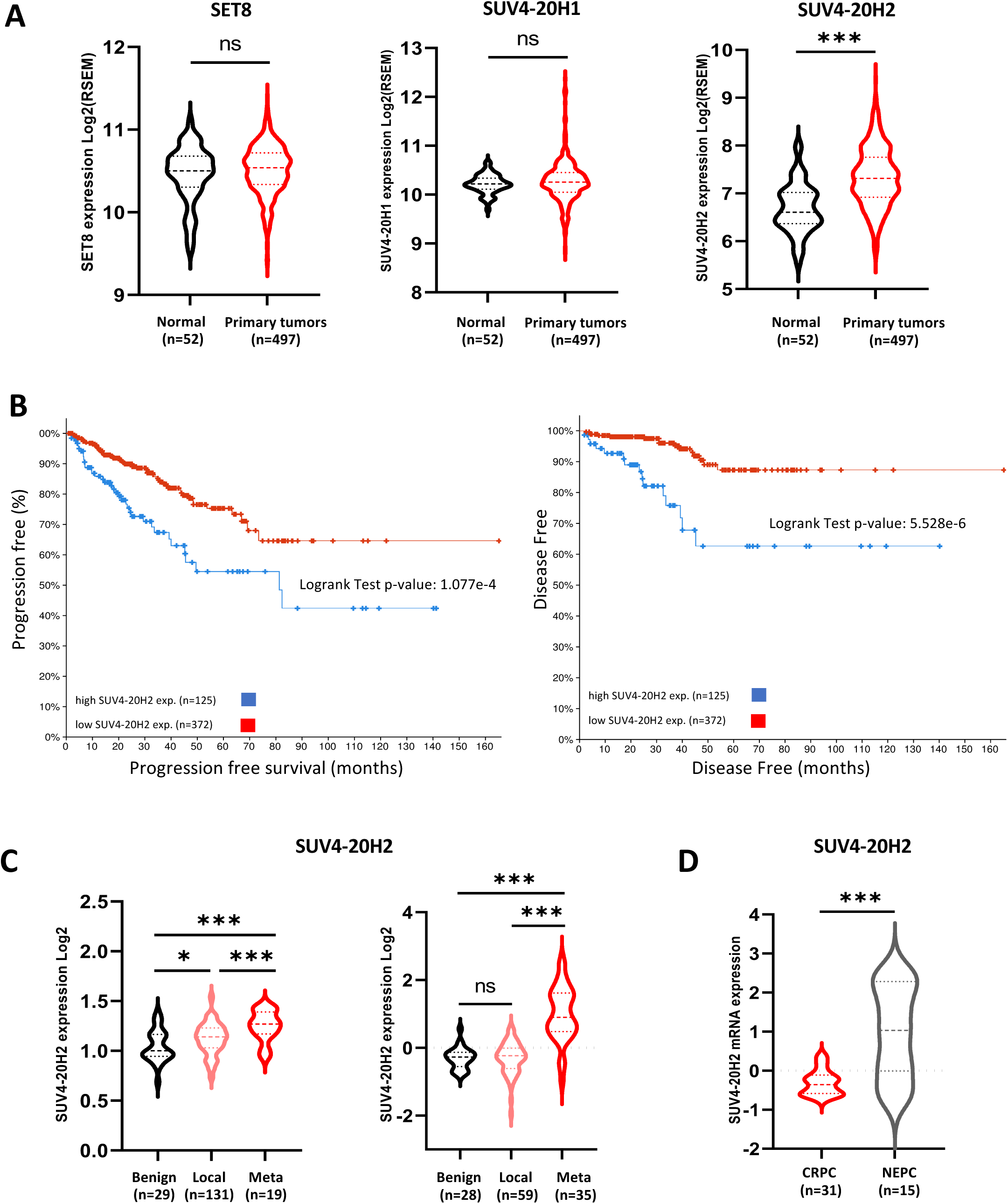
SUV4-20H2 up-regulation is a poor prognosis factor in prostate cancer. **(A)** Violin plot representing the mRNA levels of *SET8* (left), *SUV4-20H1* (middle), and *SUV4-20H2* (right) genes in normal prostate samples (52 patient samples) and primary tumor samples (497 patient samples) from the prostate TCGA database. Median (dashed line), quartile (dotted line), ns: non-significant, *** p-value < 0.0001. **(B)** Left Panel: Kaplan-Meyer analysis of progression-free interval (PFI) probability of *SUV4-20H2* expression in prostate TCGA database. The p-value is indicated on the graph. Right Panel: Kaplan-Meyer analysis of disease-free interval (DFI) probability of *SUV4-20H2* expression in prostate TCGA database. The p-value is indicated on the graph. **(C)** Violin plot representing the mRNA levels of *SUV4-20H2* in benign normal prostate samples and local or metastatic tumor samples from the GSE35988 (left) and GSE21032 (right) datasets. Median (dashed line), quartile (dotted line), ns: non-significant, **P < 0.001, ***P < 0.0001. **(D)** Violin plot showing the mRNA levels of *SUV4-20H2* (right) in castration-resistant prostate cancer CRPC tumor samples (31 patient samples) and neuroendocrine prostate cancer NEPC tumor samples (15 patient samples) from GSE21032 and GSE35988 datasets. Median (dashed line), quartile (dotted line), ns: non-significant, **p < 0.001, ***p < 0.0001.

### Targeting SUV4-20Hs induces a higher replication fork velocity that is inherently innocuous in Prostate Cancer cell lines

The elevated expression of SUV4-20H2 and its correlation with metastatic and potential aggressiveness of PCa tumors prompted us to investigate whether this enzyme is essential in prostate cancer cells. To address this question and since the functions of SUV4-20H2 overlap with SUV4-20H1 ^17^, we first utilized the selective small-molecule inhibitor A-196, which specifically targets both SUV4-20H1 and SUV4-20H2. Previous studies showed that the treatment with A196 reduces the conversion of H4K20me1 to higher H4K20me states ^27^.Consistent with these findings, a 72 hours treatment with 4 µM of A-196 was sufficient to induce the depletion of H4K20me2/3 and the concurrent increase in H4K20me1 levels in both metastatic LNCaP and DU145 prostate cell lines (Figure 2A). Despite these genome-wide epigenetic alterations, RNA sequencing revealed that the pharmacological inhibition of SUV4-20H enzymes led to relatively modest changes in gene expression levels, with only 268 genes upregulated and 26 genes slightly downregulated in metastatic castration-resistant prostate DU145 cells (Figure 2B and Supplementary Figures S3 and S4). Among them, no specific pathways or cellular processes were found to be significantly under- or over-represented.

**Figure 2:**
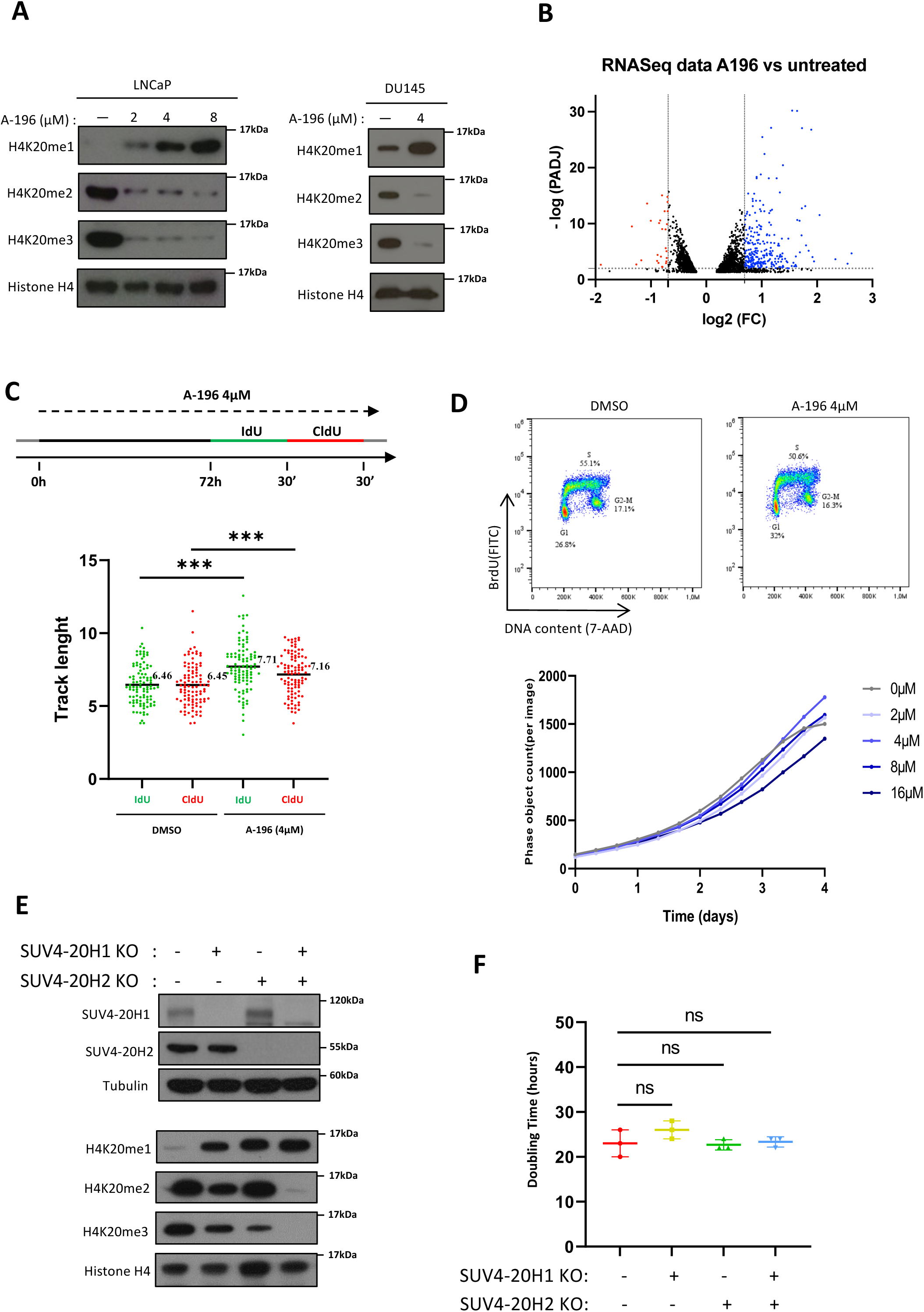
SUV4-20H inactivation induces an increase in replication fork rates in metastatic prostate cancer cells. **(A)** Immunoblot analysis showing H4K20me1, H4K20me2, and H4K20me3 levels in LNCaP and DU145 metastatic prostate cancer cell lines treated or not (DMSO) with the SUV4-20Hs pharmacological inhibitor A-196 at the indicated concentrations for 72h. Histones H3 and H4 are used as loading control, n=3. **(B)** Volcano plot representing the RNA-seq data after normalization for DU145 cell line treated or not with A-196 5μM for 10 days. Blue and red dots represent the genes that are significantly upregulated and downregulated respectively. The list of genes is shown in tables 1 and 2. FDR = 0.01 and log2(Fold Change) > 0.69 (**C**) The top diagram shows the experimental design and the timing with IdU (green) and CldU (red) incubation. Dot plot represents the IdU (green) and CldU (red)-labeled replication track lengths in DU145 cells treated or not with A-196 (4 μM) for 72h, n=3, 100 replication tracks per condition, ***p-value < 0.0001. (**D**) Upper panels are cell-cycle analysis by FACS of DU145 cells treated with A-196 (4μM) or not (DMSO) for 72h. Total DNA was stained with 7-AAD, and replicating cells was labelled with BrdU, n=3. Lower panel shows the proliferation curve of DU145 cells treated with increasing concentration of A-196 as indicated during 4 days. The number of cells was quantified as phase objective count using incucyte live cell imaging assay, n=2. **(E)** Immunoblot analysis showing SUV4-20H1, SUV4-20H2, H4K20me1, H4K20me2, and H4K20me3 levels in wild-type (WT) DU145 cells and in DU145 cells knock-out for SUV4-20H1, SUV4-20H2, or both enzymes. Histone H4 and tubulin are used as a loading control, n=3. **(F)** Graph representing the doubling time of wild-type DU145 (WT) and of DU145 cells knock-out for SUV4-20H1 or SUV4-20H2 or both enzymes (SUV4-20H1/H2 KO). Cells were counted as phase objective count using Incucyte live cell imaging assay, n=3, ns: non-significant p-value.

**Table 1:** List of downregulated genes according to the log2 (FC) value in DU145 cells after H4K20me reprogramming induced by the chemical inhibitor A196. (FDR 0.01, log2 (FC) < −0.69). The different columns correspond to Gene symbol (feature), gene ID, and log2 (FC)

**Table 2:** List of upregulated genes according to the log2 (FC) value in DU145 cells after H4K20me reprogramming induced by the chemical inhibitor A196. (FDR 0.01, log2 (FC) > 0.69). The different columns correspond to Gene symbol (feature), gene ID, and log2 (FC)

Since H4K20 methylation and SUV4-20Hs have been involved in the regulation of DNA replication ^11,14,28,29^, we next examined the rates of DNA replication forks by DNA fiber-labeling approaches. After 72 hours of treatment or not with A-196, DU145 cells were pulsed labelled with iodo-deoxyuridine (IdU) for 30 min followed by a second pulse of chlorodeoxyuridine (CldU) and the labeling replicating DNA was measured by microscopy using fluorescently labeled antibodies on DNA spread onto a glass slide. As shown in Figure 2C, replication fork rates were significantly increased by 16% above in A196-treated cells compared to control DMSO-treated cells. This higher fork velocity is likely caused by the impairment in replication origin licensing that occur upon loss of SUV4-20H enzymes ^12^. However, despite the acceleration in replication fork progression, the proliferation and the cell-cycle distribution of A-196-treated cells remained largely unaffected without the occurrence of cell death (Figure 2D).

To ascertain further these results and examine the role of each SUV4-20H enzyme in prostate cancer cells, single and double knock-out for SUV4-20H1 and SUV4-20H2 were generated in DU145 cell line using CRISPR-Cas9 approaches. As shown in Figure 2E, immunoblots with specific antibodies against SUV4-20H1 and SUV4-20H2 confirmed the successful generation of the different SUV4-20H knockout cell lines (Figure 2E, upper panels). Compared to the parental DU145^WT^ cell line, the loss of SUV4-20H1 alone resulted in a partial decrease in the levels of H4K20me2 and H4K20me3, while the loss of SUV4-20H2 mainly affected H4K20me3 (Figure 2E, lower panels). Conversely, the concomitant knock-out of both SUV4-20 enzymes resulted in a complete depletion of H4K20me2 and H4K20me3 states, as it was observed in A196 treated cells (Figure 2E). Similar to the pharmacological inhibition of these enzymes by the A-196 inhibitor, the loss of each SUV4-20H enzyme increased DNA replication fork velocity (Supplementary Figure S3) without affecting the doubling time of SUV4-20H-inactivated cells (Figure 2F). Altogether, these results confirm the efficacy of A-196 to inhibit SUV4-20H enzymes and show that the lack of conversion of H4K20me1 to higher H4K20me states leads to an increased rate of replication fork progression, which remains inherently non-detrimental to the proliferation and survival of prostate cancer cells.

### The pharmacological inhibition of SUV4-20H enzymes induces a lethal synergy in combination with TOP2 poisons, notably etoposide, in prostate cancer cells

Given the higher velocity of DNA replication forks upon SUV4-20H inhibition, we asked whether prostate cancer cells treated with A-196 could display enhanced sensitivity to specific inducers of replication stress, notably those commonly used in cancer treatment. Following a 72 hours A-196 treatment in DU145 cell line to induce H4K20me reprogramming, we evaluated the interaction of A-196 with six selected drugs using a concentration matrix test. This involved the assessment of increasing concentrations of each single drug with all possible combinations of the second drug. For each combination sample, the cell viability was quantified by measuring the percentage of live and dead cells with the Celigo image cytometer. The results are shown in Figure 3A according to the combination and efficiency index calculated for each drug combination. Strikingly, a positive interaction was observed between A-196 and TOP2 poisons with the highest combination and efficiency index achieved with etoposide (Figure 3A). Remarkably, the combination of A-196 with etoposide exhibited a lethal synergy in all PCa cell lines, with the strongest synergy with the metastatic castration-resistant DU145 and PC3 cell lines (Supplementary Figure S4A). Long-term colony formation assays further confirmed a synthetic lethality between A-196 and sub-lethal doses of etoposide in different prostate cancer lines (Figures 3B and 3C). Remarkably, the lethal interaction between A196 and of etoposide was more effective than this previously found with the EzH2-mediated H3K27me3-targeting inhibitor GSK126, which showed predominantly additive effects in our experimental conditions (Supplementary Figure S4B). Collectively, these data demonstrate an unique lethal synergy between the chemical inhibitor A-196 and TOP2 poisons, notably etoposide, in all metastatic prostate cancer cell lines tested.

**Figure 3:**
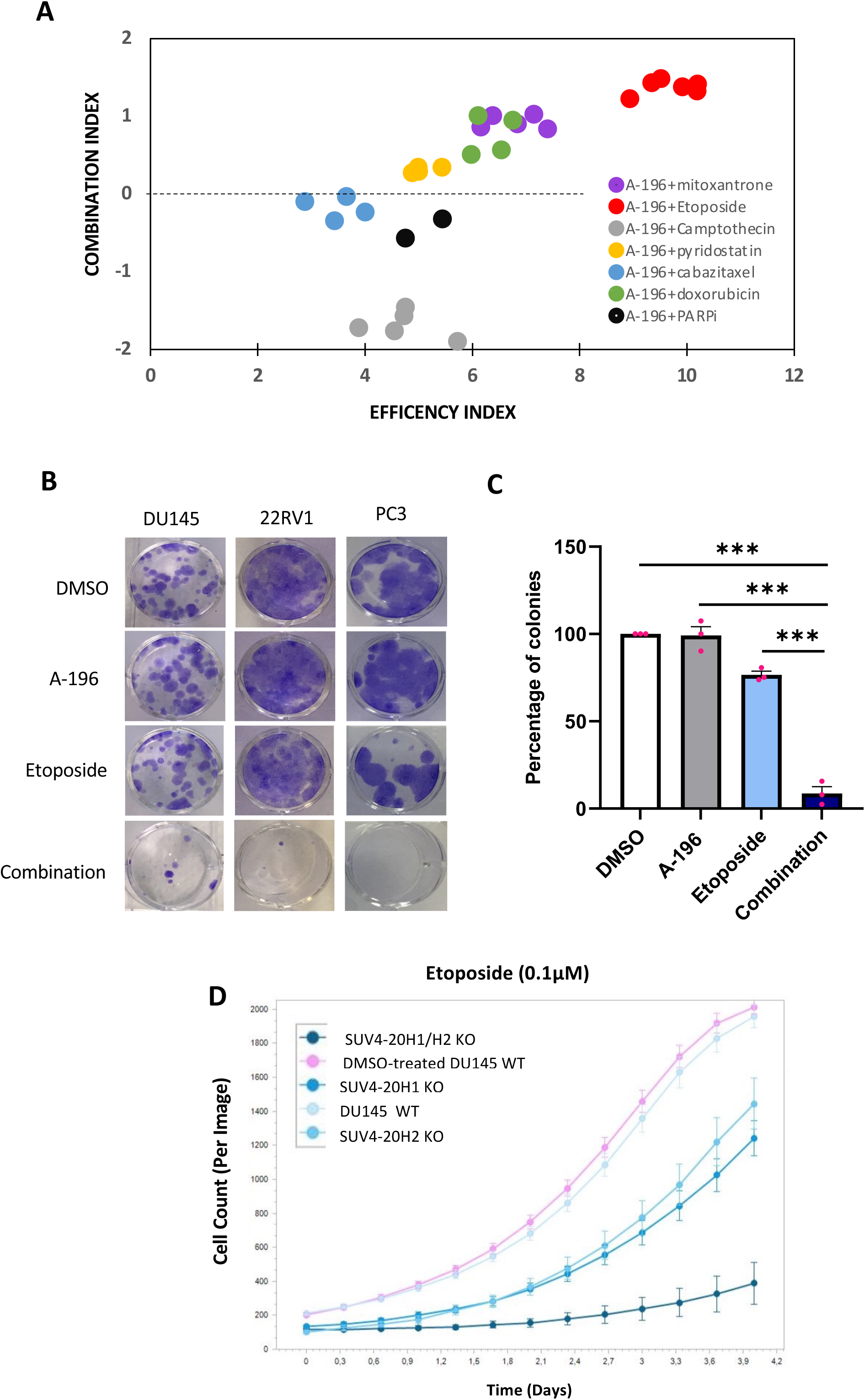
targeting SUV4-20Hs enzymes induces a lethal synergy specifically in combination with etoposide. **(A)** Scatter plot representing the combination index (y) and the efficiency index (x) of A196 combination with different replication stress inducers during 72 hours in metastatic prostate cancer cell lines. Each dot represents the results of a combination experiment. Synergistic effects are observed when the combination index is upper 0 (dashed bar). **(B)** Representative images of clonogenic assays for DU145, 22RV1, and PC3 cell lines treated with A-196 (4μM) and various sub-lethal concentrations of etoposide (0.2, 0.63, and 0.3 μM) for 72h, as indicated. **(C)** Bar plot revealing the percentage of colonies for DU145 cell lines treated with A-196 4μM and etoposide 0.2μM for 72h as indicated, n=3, ***p < 0.0001. **(D)** Graph representing the proliferation rate of wild-type DU145 and DU145 cells depleted for SUV4-20H enzymes, as indicated, and treated with etoposide (0.1μM) for 4 days. Cells were counted as phase objective count using incucyte live-cell imaging assay, n=4.

### The synthetic lethality with etoposide requires the inactivation of both SUV4-20H1 and SUV4-20H2

To ascertain whether the enhanced sensitivity of prostate cancer (PCa) cells to etoposide was linked to the inhibition of either SUV4-20H1 or SUV4-20H2 or both enzymes, we investigated the sensitivity of the different SUV4-20H knock-out DU145 cell lines to a sub-lethal dose (0.1 µM) of etoposide over four days using the incucyte live-cell imaging system. The results are shown in Figure 3D. As anticipated, the growth of the parental cell line upon 0.1 µM of etoposide was comparable to control DMSO-treated cells. In contrast, the proliferation of cells depleted for each enzyme was partially reduced and virtually abolished when both enzymes were depleted (Figure 3D). These results demonstrate that the sensitivity of DU145 cells to etoposide depends on each SUV4-20H enzyme and is maximized when both SUV4-20H enzymes are inactivated.

### SUV4-20H inhibition synergistically enhances the chromatin accumulation of TOP2 cleavage complexes and DNA breaks upon etoposide

Compared to treatments with each drug, flow cytometry analysis showed that the combination of etoposide and A-196 resulted in the accumulation of cells in late S and G2 phases (Figures 4A). This cell-cycle arrest was accompanied by increased cell death in a dose-dependent manner (Figure 4B). It is known that etoposide acts by stabilizing TOP2 cleavage complexes (TOP2ccs), notably behind the replication forks, and promoting DSBs via the inhibition of TOP2-mediated DNA re-ligation stage^30^. To investigate the underlying mechanisms of increased PCa cell sensitivity to etoposide upon SUV4-20Hs inactivation, the levels of TOP2α subunit covalently bound to chromatin were quantified before and after A-196 treatment using a heparin-based extraction protocol coupled with immunoblotting ^31^. Heparin treatment extracts TOP2 molecules that are not covalently attached to DNA into a soluble fraction, enabling quantification of TOP2ccs by evaluating the levels of TOP2α in the insoluble fraction. DU145 cells were incubated or not with A-196 for 72 hours followed by 3 hours treatment with increasing doses of etoposide. The levels of TOP2α in soluble and insoluble fractions were then examined by immunoblotting after heparin extraction. While the steady-state levels of TOP2α remained constant across all tested conditions before heparin extraction, the levels of TOP2ccs induced by etoposide were significantly higher in A-196 treated cells than in controls and, importantly, exponentially increased in an etoposide dose-dependent manner (Figures 4C-4D). These results indicate that SUV4-20H inhibition potentiates the cytotoxicity of etoposide by enhancing the trapping of TOP2ccs complex on chromatin.

**Figure 4:**
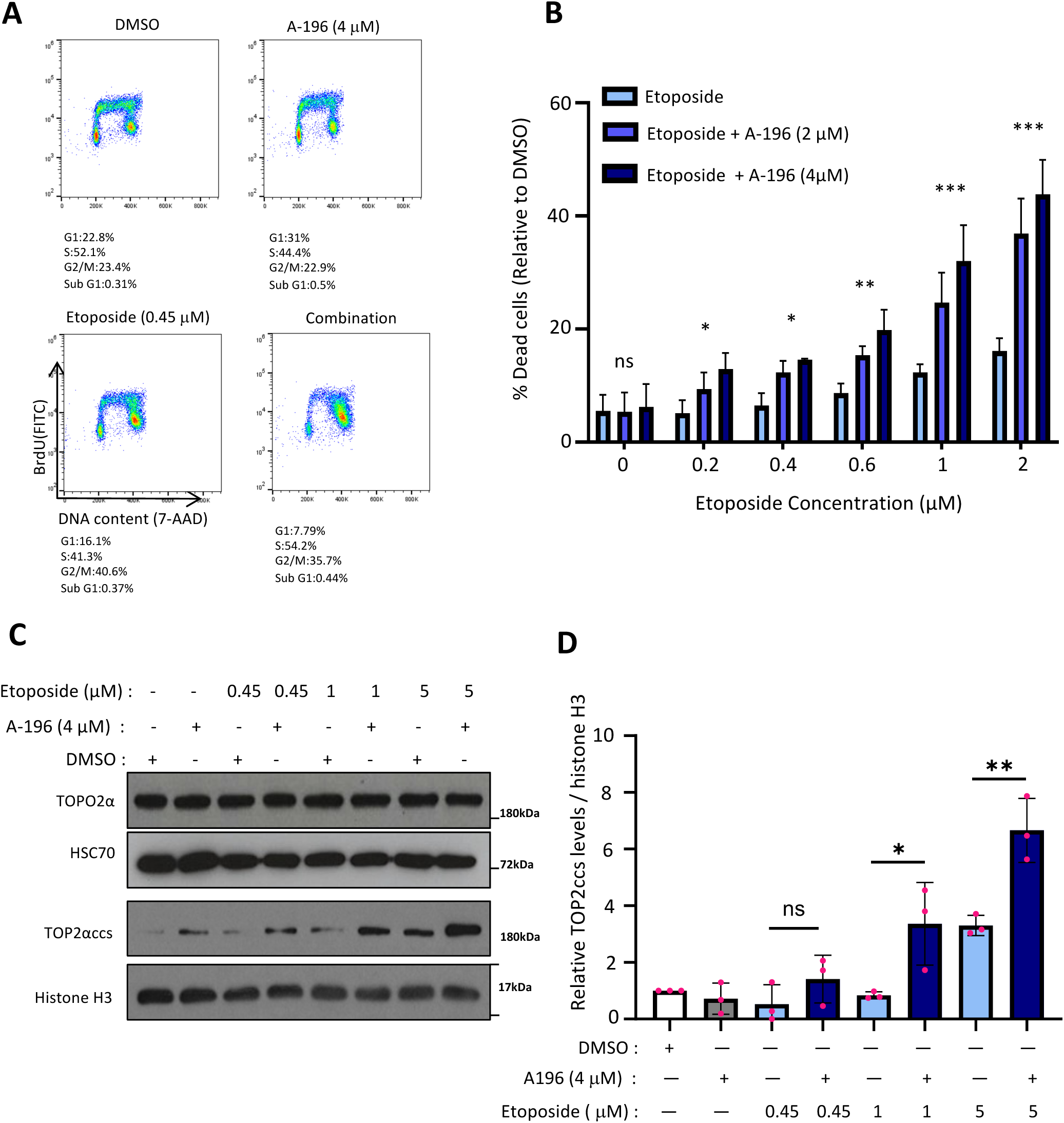
the inhibition of SUV4-20Hs improves the accumulation of TOP2ccs complexes on chromatin upon etoposide. **(A)** Cell cycle distribution of DU145 cells treated with DMSO or A-196 (4μM) for 72h and then for 24h with etoposide (0.45μM) as indicated. Total DNA was stained with 7-AAD, and nascent DNA was labeled with BrdU, n=3. (B) Bar plot quantification of dead cells measured by propidium iodide staining after treatment with A-196 and etoposide as indicated. ** p-value < 0.01, *** p-value < 0.001. **(C)** Immunoblot analysis showing the total levels of TOP2α (upper panel) and of TOP2αccs (lower panel) in DU145 cells treated with DMSO or A-196 (4μM) for 72h and then treated with increasing doses of etoposide for 3 hours as indicated. HSC70 is used as loading control in total cell extracts and histone H3 is used as quantitative control of chromatin-bound proteins after heparin extraction, n=3. **(D)** Bar plot quantification of TOP2αccs levels in control and treated cells as indicated above using ImageJ software, n=3, * p-value ≤ 0.05, ** p-value < 0.01, *** p-value < 0.001.

To examine whether the extensive etoposide-induced trapping of TOP2ccs upon SUV4-20H inhibition was followed by higher levels of DNA damage, the levels of double-strand breaks (DSBs) were measured by pulse-field gel electrophoresis using the same treatment protocol than in Figure 4. The results are shown in Figures 5A and 5B, with untreated or DMSO-treated cells serving as negative controls and cells treated with a high concentration of etoposide (10 μM) as a positive control for DSBs. While A-196 treatment alone that did noy induce DNA breaks, etoposide triggers DSBs in a dose-dependent manner in control cells as expected (Figures 5A and 5B). Upon A-196 treatment, the levels of DSBs induced by etoposide were significantly enhanced (Figures 5A and 5B). To further substantiate that the lack of SUV4-20H activity strongly increases DSB formation in response to etoposide, we examined the focal accumulation of 53BP1 and of the phosphorylated form of the histone variant H2A.X (γ-H2AX) by immunofluorescence. These markers are well established indicators of DNA breaks with known dynamics in DNA damage responses^32^. Following a 72-hour treatment with or without A-196, a sub-lethal concentration of etoposide (0.1 μM) was added for 24 hours, and cells were subsequently stained with specific antibodies for 53BP1 and γ-H2AX. Representative images and quantification of the results are shown in Figures 5C and 5D, respectively. Consistent with the pulse-field gel electrophoresis results (Figures 5A and 5B), treatment with etoposide induced a significantly higher percentage of 53BP1 and γ-H2AX positive cells when cells were pre-treated with A196 (Figures 5C and 5D). Furthermore, 53BP1 and γ-H2AX immunostaining appeared as larger and more intense foci in nuclei treated with the drug combination than in nuclei treated only with etoposide (Figure 5C). Similar results were obtained in SUV4-20H1 and SUV4-20H2 knock-out DU145 cells (Supplementary Figure S5). Collectively, these findings demonstrate that the inhibition of SUV4-20H enzymes and likely the consequent alterations in H4K20me levels create a chromatin environment conducive to the enhanced trapping of TOP2 cleavage complexes by etoposide. This leads to increase accumulation of DNA double strand breaks, resulting in S/G2 cell-cycle arrest and massive cell death.

**Figure 5:**
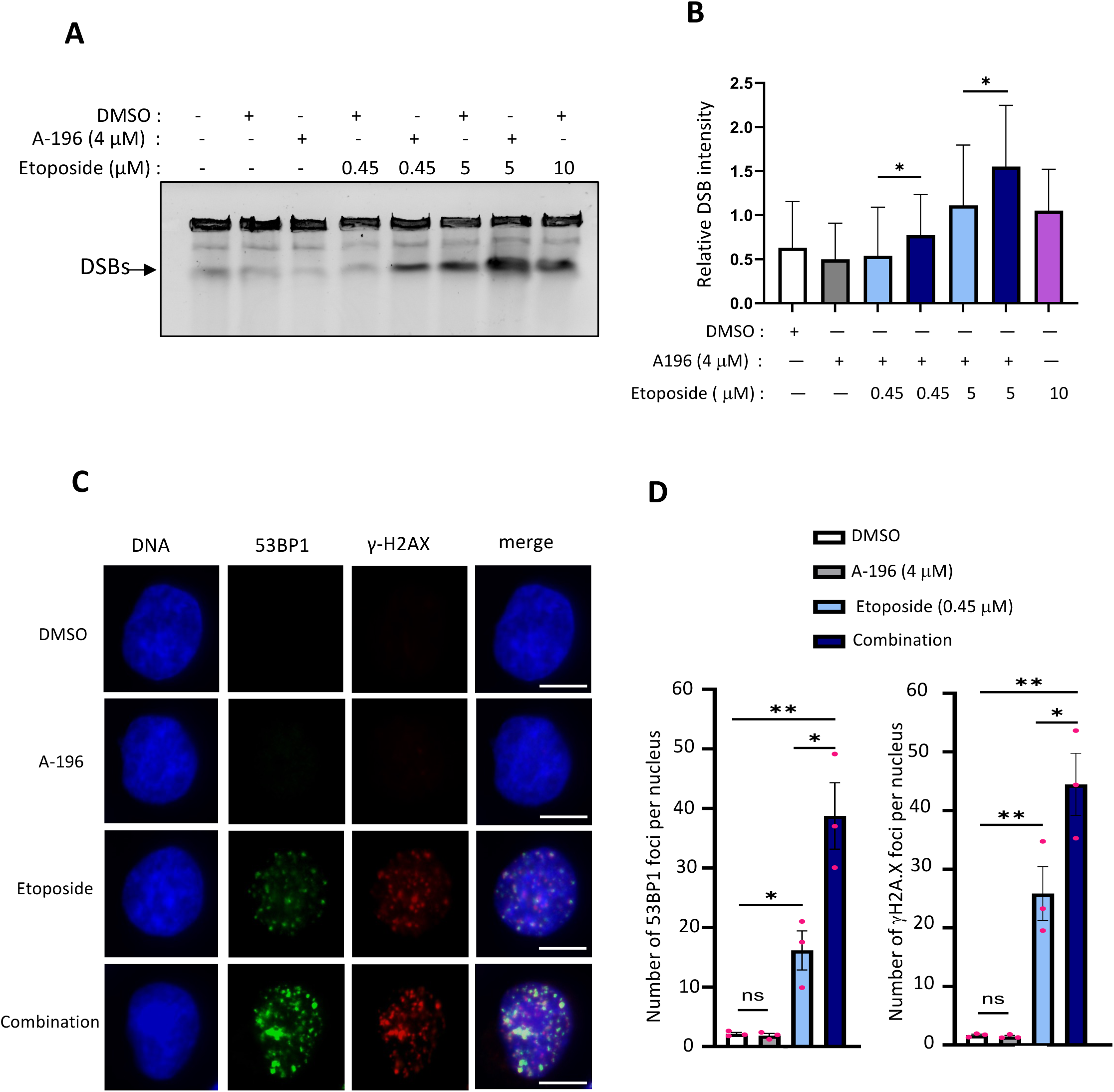
The enhanced toxicity of etoposide in A-196 treated cells is due to an increase in DSB formation. (A) Representative image of ethidium bromide staining of uncut DNA (upper band) and double-strand DNA breaks (DSBs) (lower band) separated by pulse-field gel electrophoresis in DU145 cells treated with DMSO or A-196 (4μM) for 72 hours before treatment for 3 hours with increasing doses of etoposide as indicated. **(D)** Bar plot representing the intensity of DSBs formed in control and treated cells as indicated, n=4, *P < 0.01. **(C)** Representative images of 53BP1 and γ-H2AX staining in DU145 cell line treated with DMSO or A-196 (4μM) for 72h, before incubation with etoposide (0.45 μM) for 24 hours. Scale bar = 10μm. **(D)** Bar plots representing the number of 53BP1 and γ-H2AX foci per nucleus of DU145 cell line treated or not as indicated, n=3, mean of each experiment (pink dot), >200 cells per condition, ns: non-significant, *p-value < 0.01, **p-value < 0.001, ***p-value < 0.0001.

### The repair of TOP2-mediated DSBs by Homologous Recombination is hindered by SUV4-20H inhibition

We observed that DSBs induced by the combination of A-196 and etoposide were accompanied by a pronounced activation of the DNA-damage response kinases ATM and DNA-PKcs, and to a lesser extend of ATR, as evidenced by increased levels of the phosphorylation status of these three kinases (Figure 6A). Strikingly, this activation was accompanied by a significant phosphorylation of the RPA32 subunit (Figure 6A) and the nuclear accumulation of RPA70 subunit (Figure 6B), thereby indicating an activation of the Homologous Recombination (HR) pathway and the formation of single-strand DNA at DNA damage sites. The focal accumulation RPA normally involves its rapid replacement with RAD51 to form a nucleofilament competent for HR ^33^. However, we observed lower levels of RAD51 nuclear foci in A-196 treated cells compared to control cells following etoposide treatment (Figure 6C). This suggests that the switch from RPA to RAD51 at DNA breaks might be somehow compromised by the inhibition of SUV4-20H activity, potentially resulting in inefficient repair of TOP2-mediated DSBs. To further explore this hypothesis, DU145 cell were pre-treated for three 72 hours with A196 in order to reprogram the levels of H4K20 methylation states. Subsequently, cells were exposed to 0.45 μM of etoposide for 24 hours to induce quantitative levels DSBs. After washing, control and A-196 treated cells were cultured with etoposide-free medium and the ability of cells to repair their DNA was evaluated by monitoring the levels of RPA, RAD51, 53BP1 and γH2A.X foci after 6 hours, 24 hours and 48 hours. In control cells treated with etoposide alone, the transition from RPA to RAD51 foci led to a rapid decrease of RPA and RAD51 foci to background levels was observed 24 hours after etoposide removal (Figures 6D and 6E). Consistently, 53BP1 and γH2A.X focal accumulation followed similar kinetics, indicating an effective repair of DSBs (Figures 6F and 6G). In contrast, A-196 treated DU145 cells displayed defects in RAD51 focal accumulation that peaked only 6 hours after etoposide removal (Figure 6E). This coincided with a persistence of high levels of RPA and DNA PKCS phosphorylation (supplementary Figure S6). Accordingly, the levels of RPA, 53BP1 and γH2A.X focal accumulation remained elevated and showed only a slow reduction over 48 hours compared to control cells (Figure 6D and 6E). Noted of, similar results were obtained in the prostate castration-resistant 22RV1 cell line, indicating that they were not specific to a particular prostate cell line (Supplementary Figure S7). Moreover, FACS analysis revealed that a 24-hours pulse of sub-lethal concentration of etoposide in A196-treated cells was sufficient to induce a S/G2 cell cycle arrest followed by cell death as shown by the appearance of a sub-G1 fraction (supplementary figure S8). Thus, the inhibition of SUV4-20H enhances the potency of etoposide not only by increasing TOP2ccs trapping but also by impairing the switch from RPA to RAD51, which disrupts the repair of TOP2-induced DNA breaks.

**Figure 6:**
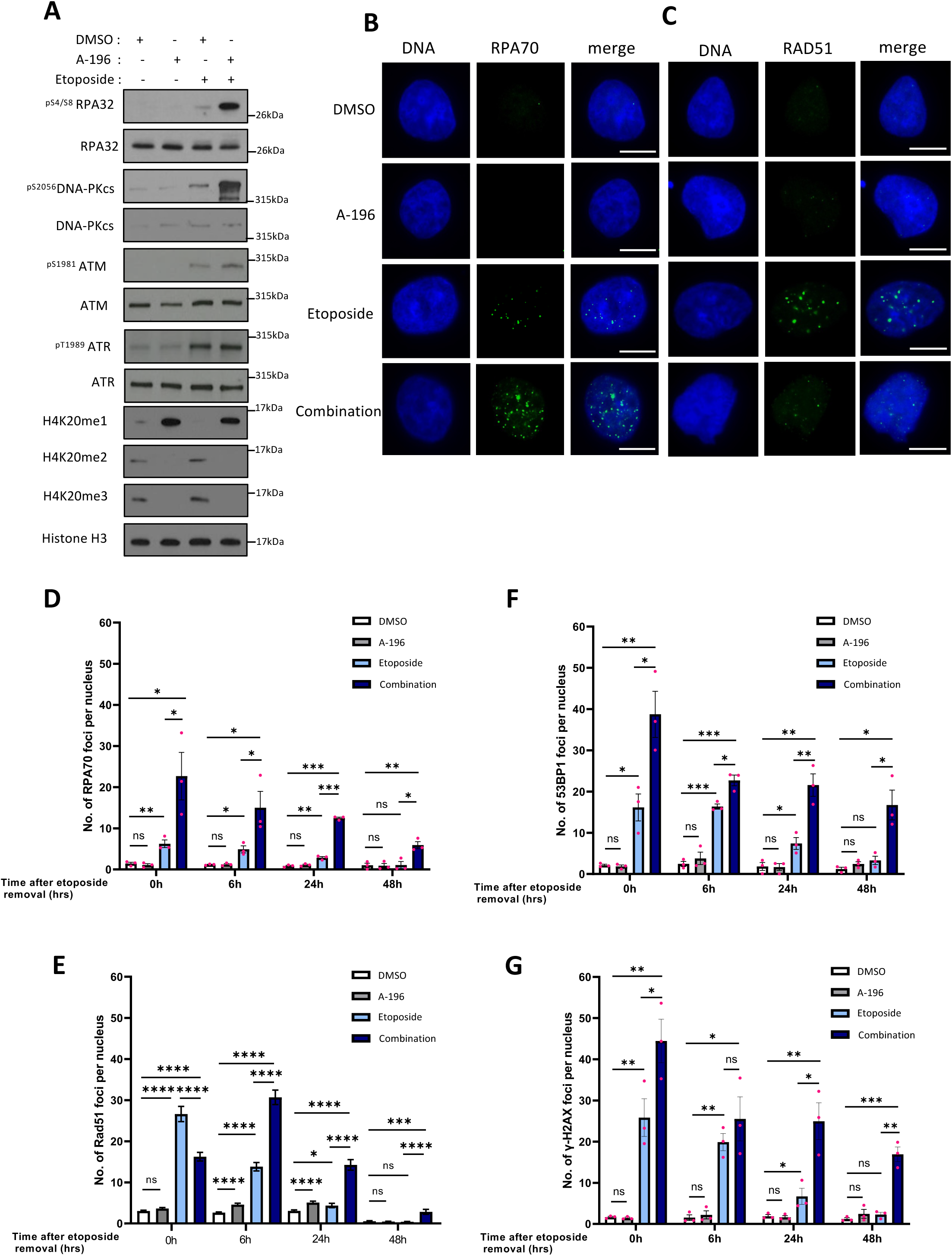
A-196 treatment impairs the ability of cancerous cells to repair TOP2-induced DNA breaks. **(A)** Immunoblot analysis showing the expression levels of indicated proteins in DU145 cell line treated with DMSO or A-196 (4μM) for 72 hours, and then exposed to etoposide (0.45μM) for 24 hours as indicated. n=3. **(B)** Representative images of RPA70 staining in DU145 cells treated as above. Scale bar = 10μm. **(C)** RAD51 staining in DU145 cells treated as above. Scale bar = 10μm. **(D)** Bar plot representing the number of RPA70 foci per nucleus of DU145 cells treated with DMSO or with A-196 (4μM) for 72h before treatment with etoposide (0.45μM) for 24h and then collected at different times after etoposide removal as indicated, n=3, mean of each experiment (pink dot), >200 cells per condition, ns: non-significant, *P < 0.01, **P < 0.001, ***P < 0.0001. **(E)** Bar plot representing the number of RAD51 foci per nucleus of DU145 cells treated as above. =3, mean of each experiment (pink dot), >200 cells per condition, ns: non-significant, *P < 0.01, **P < 0.001, ***P < 0.0001 (**F**) Bar plot representing the number of R53BP1 foci per nucleus of DU145 cells treated as above. =3, mean of each experiment (pink dot), >200 cells per condition, ns: non-significant, *P < 0.01, **P < 0.001, ***P < 0.0001. **(G)** Bar plot representing the number of γ-H2AX foci per nucleus of DU145 cells treated as above. =3, mean of each experiment (pink dot), >200 cells per condition, ns: non-significant, *P < 0.01, **P < 0.001, ***P < 0.0001.

### TOP2 poisoning combined with SUV4-20H inhibition leads to enhanced anti-tumor activity *in vivo*

Our *in vitro* results suggest that targeting SUV4-20H enzymes is a potential efficient strategy to increase the efficiency of etoposide and eventually reduce its effective dosage in the clinic to lower side effects. To validate this hypothesis and demonstrate *in vivo* the enhanced anti-tumor activity of etoposide when SUV4-20H are inhibited, mice bearing 100 mm^3^ DU145 prostate tumors were treated twice a week with vehicle (20% DMSO + corn oil), etoposide (2.5 mg per kg), A-196 (7.5 mg per kg), or the combination of the two drugs, as illustrated in Figure 7A. Importantly, no toxicity was observed in all animal groups as shown by body weight measurement (Figure 7B). Moreover, As shown in Figures 7C and 7D, monotherapy with A-196 or etoposide did not reduce significantly tumor growth, indicating that these two drugs are innocuous in vivo at these concentrations. In contrast, the combination significantly delayed tumor growth (Figures 7C and 7D). To further support these results, mice were xenografted with wild-type or SUV4-20H1 and SUV4-20H2 double knock-out DU145 cells. Once tumors reached 100 mm3, they were treated with the vehicle (DMSO) or etoposide (2.5 mg or 6 mg per kg) twice a week. Of note, without any treatment, tumors derived from wild-type DU145 grew slightly slower than those derived DU145 inactive for SUV4-20H enzymes (Figure 7E). Upon 6 mg per kg of etoposide, the growth of double SUV4-20H knock-out tumors was strongly impaired, and even regressed. In contrast, no difference was observed for wild-type tumors (Figures 7E and 7F). Importantly, these effects were observed in the absence of toxicity for the animals (Figure 7G). Taken together, our results demonstrate that targeting of SUV4-20H1/H2 enzymes and reprogramming H4K20 methylation can strongly improve *in vivo* response to etoposide while reducing its effective dose in malignant tumors and without toxicity for the animal health.

**Figure 7:**
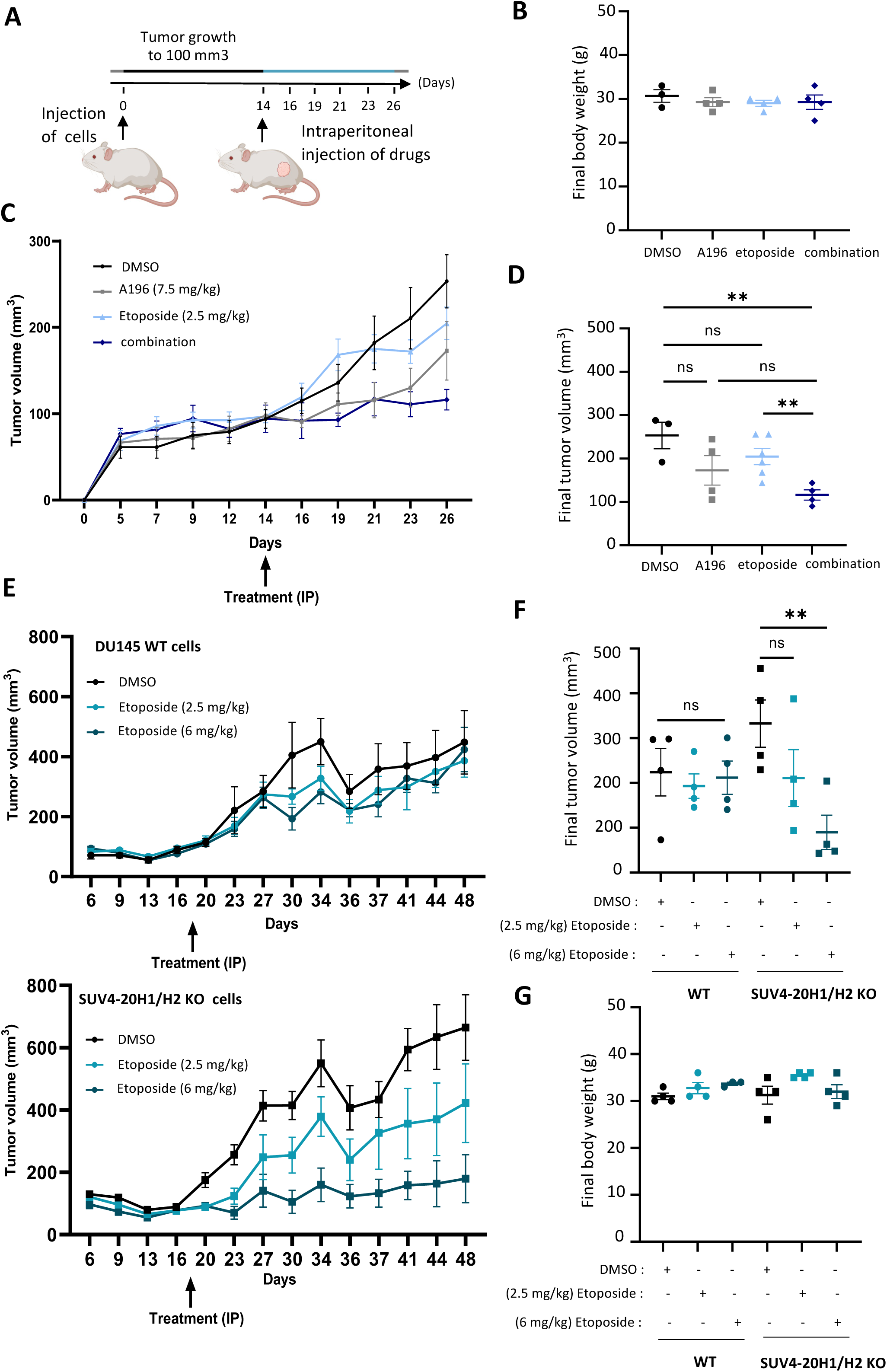
The depletion of SUV4-20H enzymes improves *in vivo* therapeutic response to etoposide. **(A)** Schematic representation of experimental procedure and drug administration. **(B)** Graphical representation of the body weight measurement of mice after three weeks of treatment as indicated. Intact athymic nude males were inoculated with DU145 cells by subcutaneous injection for 2 weeks. All mice were divided then into four cohorts of 8 animals: Vehicle (DMSO+cornoil), A-196(7.5 mg/kg) etoposide (2.5 mg/kg), and combination. Tumor sizes were measured every 3 days. **(C)** Tumor growth curves of the DU145-derived xenograft tumors as indicated. **(D)** Graphical representation of the tumor growth volume of treated mice as indicated after three weeks of treatment. ns: non-significant, ** p-value < 0.001. **(E)** Growth curves of wild-type (WT) DU145 xenograft tumors (upper panel) and SUV4-20hs knock-out DU145 xenografts tumors (lower panel), two weeks after subcutaneous DU145 cell injection in mice treated (IP injection) with DMSO or with etoposide as indicated. Tumor sizes were mainly measured every 3 days. **(F)** Graphical representation of the final volume wild-type and SUV4-20H depleted DU145 xenograft tumors in mice treated with etoposide every three days (IP injection at two different concentrations as indicated). ns: p-value > 0.05, **p-value < 0.001. **(G)** Graphical representation of the final body weight of mice treated as above.

## DISCUSSION

Despite intensive multimodal therapy, Prostate Cancer (PCa) remains a major cause of death due to the largely incurable nature of its metastatic and castration-resistant stages^34^. It is therefore crucial to identify new markers for patients with the highest risk for developing aggressive disease and to propose new therapeutic options. In this study, we present evidence for the first time that the expression levels of histone H4K20 methyltransferase SUV4-20H2 can serve as a new marker for aggressive prostate cancer independently to the Gleason score (Figure 1). Consistent with this, elevated levels of SUV4-20H2 are associated with a poor prognosis and its gradual increase during disease progression correlates with the risk of metastasis (Figure 1). Using genetic and pharmacological approaches, we also demonstrate that targeting SUV4-20H2 and its paralog SUV4-20H1 leads to a global reprograming of H4-K20 methylation states, resulting in higher replication fork velocity (Figure 2). Remarkably, these chromatin and replication alterations do not induce overt toxicity in prostate cancer cell lines, but create conditions for synthetic lethality when combined with sub-lethal concentration of topoisomerase-II (TOP2) poisons, such as etoposide (figure 3). Conspicuously, we reveal that inhibiting SUV4-20H enzymes significantly enhances the potency of etoposide by increasing the trapping of TOP2 cleavage complexes on chromatin, thereby inducing DNA breaks and reducing the capacity of cells to repair these breaks through homologous recombination (Figures 4-6). Finally, using xenografted mice models, we demonstrate that the pharmacological inhibition of SUV4-20Hs enzymes or their genetic deletion are indeed sufficient to enhance the *in vivo* antitumor activity of etoposide while reducing its dosage at non-toxic levels (Figure 7). Altogether, this holds a potential new strategy for improving the effectiveness of etoposide in the treatment of advanced prostate cancer stages.

The progressive increase in SUV4-20H2 levels as the disease advances is not unique to prostate cancer. This has been also reported in hepatocellular carcinoma, clear renal cell carcinoma and pancreatic cancer^23,35,36^, indicating it may be a specific epigenetic feature of certain solid tumors. The mechanisms leading to SUV4-20H upregulation in cancer cells remain unclear. In pancreatic cancer, this could be partially due to gene amplification^35^. There is no evidence for this in prostate tumors, suggesting that is the consequence of a broader biological mechanism. It is also remains uncertain whether SUV4-20H2 upregulation impacts prostate cell behavior and whether it is associated with specific changes in H4K20 methylation states. Paradoxically, immunohistochemistry experiments have suggested that prostate tumors often exhibit low levels in H4K20me1 and H4K20me2 while the levels H4K20me3, primarily mediated by SUV4-20H2, remains unchanged^24^. However, further analysis with next-generation antibodies specific to each H4K20me state is needed to confirm these initial results and to determine whether H4K20me alterations in prostate tumors are related to SUV4-20H2 upregulation.

The absence of apparent cellular toxicity upon the pharmacological inhibition of SUV4-20H enzymes is consistent with previous results in various human cell lines^20,37^. By generating single and double knock-outs for SUV4-20H2 and SUV4-20H1, we confirmed that the loss of SUV4-20H enzymes is largely innocuous for prostate cancer cells and likely not key drivers in cancer cell proliferation and survival *in vitro*. We noticed, however, higher replication fork speed when SUV4-20H enzymes were inhibited or inactivated in prostate DU145 cells. Since H4K20me2 and H4K20me3 contributes to the licensing of replication origins by serving as binding sites for ORC complexes ^11,12,38^, increasing fork velocity may be a compensatory mechanism used by cancer cells to offset reduced origin activity upon loss of SUV4-20H activity^39^.

Interestingly, recent evidences suggest this may lead to increased genomic instability^40^, at least when fork speed exceeds a certain threshold, which may vary depending on the nature and physiological state of the cell. This could in part explain why prostate cancer cells deficient for SUV4-20H activity are particularly sensitive to certain specific replication stress inducers, leading to the accumulation of the prostate cancer cells at the end of S phase followed by massive cell death.

In hepatocellular carcinoma, it has been also recently reported that the inhibition of SUV4-20H2 with the chemical inhibitor A-196 sensitizes cells to the PARP inhibitor Olaparib^23^. Our screening results did not confirm such a synergistic effect in prostate cancer cells. In contrast, our *in vitro* and *in vivo* results show that the inhibition or deletion of SUV4-20H enzymes confers a higher sensitivity of prostate cancer cells to the TOP2-poison etoposide. Mechanistically, we demonstrated that the loss of SUV4-20H enhances the etoposide-mediated accumulation of trapped TOP2 complexes and the levels TOP2-induced DNA breaks. This is likely related to a better accessibility of TOP2 complexes to etoposide poisoning in absence of SUV4-20H activity on H4K20me1 in chromatin. Indeed, it has been recently shown that the loss of H4K20me2/3 and the accumulation of H4K20me1 facilitates chromatin openness and accessibility by disrupting chromatin folding throughout the cell cycle^41^. Of note, the treatment with inhibitors of histone deacetylase (HDAC) also induces chromatin decompaction and the accumulation of TOP2ccs complexes following treatment with TOP2 poisons^4,42^. However, in contrast to HDAC inhibitors that induces massive transcriptional alterations, our data show that the treatment with the SUV4-20H inhibitor A-196 has only a mild impact on gene expression. Thus, loosening-up the chromatin structure upon SUV4-20H inhibition does not globally affect gene regulation or specifically occurs in gene-poor chromatin regions. Compared to the clinic use of HDAC, this is likely unique and clearly represents an advantage to limit off-target and adverse secondary effects in combination with etoposide. This is clearly of paramount interest, as the dosage reduction of etoposide is an important issue due to its relative toxicity in the clinic, particularly in elderly patients with advanced prostate cancer.

In addition to its effects on the trapping of TOP2 complexes and DNA damage following etoposide treatment, another feature of A196 treatment is to hinder the repair of TOP2-mediated DNA breaks by delaying the switch from RPA to RAD51 at damaged DNA sites. Interestingly, it was recently shown that the catalytic domain of SUV4-20H2 can interact with RAD51^23^, suggesting a potential direct role of SUV4-20H in homologous recombination mechanisms. However, defects in homologous recombination following the loss of SUV4-20H are also linked to alterations in H4K20 methylation states. Indeed, we observed that the impairment in the repair of etoposide-induced DNA breaks occurs only when changes in H4K20 methylation, triggered by SUV4-20H inhibition, precede etoposide treatment. In this regard, it was reported that higher levels in H4K20me1 are sufficient to recruit 53BP1^43^ even in the absence of H4K20me2, thereby limiting homologous recombination by antagonizing the DNA end resection activity of BRCA1-BARD1 and the subsequent loading of RAD51 on damaged DNA. Consistent with this, we show that A-196 treatment in prostate cancer cells has not impact on 53BP1 recruitment, which is even increased after etoposide treatment.

The combination of A-196 with etoposide was previously tested in U2OS cells where only a mild sensitivity was reported ^37^, suggesting that the lethal synergy between these drugs might be specific to prostate tumors. This might be dependent on the up-regulation of SUV4-20H2 in prostate cancer cells, but also to the high levels of TOP2 in this cancer where it contributes to disease progression and resistance to anti-androgen therapy^44,45^. The elevated levels of TOP2 have suggested a potential benefit of targeting it in this disease. Thus, the TOP2-poisons mitoxantrone and etoposide have already been included as standard of care treatment for metastatic and castration-resistant prostate cancer^6^. However, given the inconvenient risk-benefit ratio, these drugs are currently limited to combination with platinum agents to treat the most aggressive forms of neuroendocrine prostate cancer^6,45^. As an alternative strategy, the combination therapy using an inhibitor of SUV4-20Hs with etoposide could be utmost interest in clinic for managing advanced and metastatic prostate cancer. This will however require to understand further SUV4-20H enzyme functions in both normal and prostate cancerous cells. Notably it will be essential to determine more precisely where these enzymes bind to chromatin, what are their substrates beyond histone H4K20, and how they are functionally linked to topoisomerase activity during the cell cycle. From a clinical point of view, it will also be necessary to identify new inhibitors which are more effective *in vivo* and which could target each SUV4-20H enzyme. The optimization of drug delivery within the prostate tumor and associated metastases will be also critical for future use of these drug combinations in clinical trials.

## MATERIALS & METHODS

### Analysis of prostate cancer patient data

Gene expression values from the Prostate Cancer and PanCancer TCGA cohorts were retrieved from the Firebrowse website reporting processed TCGA data (http://firebrowse.org)^25,46^. Expression values from normal (N=52) and prostate cancer (N=497) specimen were determined by RNA sequencing and expressed as mRNA gene expression. They were quantified using the RSEM package; https://github.com/deweylab/RSEM). Genomic and clinical data analyzed are freely available, re-usable and anonymized, and were retrieved from the cBioPortal for Cancer Genomics portal. The following detailed parameters were extracted: patient age, recurrent structural variants and mutations, neoplasm disease lymph node stage, Gleason score, fraction genome altered, MSI MANTIS score, aneuploidy score, Buffa and Winter hypoxia score. Gene expression values from GSE21032 ^47^ and GSE35988 ^48^ datasets were collected from the GEO repository (https://www.ncbi.nlm.nih.gov/ gds) to assess gene expression changes in primary, local and metastatic prostate tumors.

### Cell culture

Prostate cancer cell lines were purchased from ATCC. DU145 cell lines were cultured in Dulbecco’s modified Eagle’s medium (DMEM) + Glutamax (Gibco). LnCaP, 22RV1, and PC3 cell lines were cultured in Roswell Park Memorial Institute medium (RPMI) + Glutamax (Gibco). Culture medium was supplemented with 10% fetal bovine serum and 1% penicillin/streptomycin. Cells were incubated at 37°C, 5% CO2.

### Drugs and antibiotics

The following reagents were used: A-196 (MedChemexpress HY-100201), Etoposide (Sigma-Aldrich E1383), Camptothecin (Sigma-Aldrich C9911), Mitoxantrone (Sigma-Aldrich M6545), Doxorubicin (Sigma Aldrich D1515), Pyridostatin (MedChem express HY-15176A), Cazabitaxel (Sigma Aldrich SML2487), Olaparib (MedCemexpress HY-10162), Puromycin (InvivoGen).

### Crispr/Cas9-mediated SUV4-20H1 and SUV4-20H2 KO models

Knock-out of SUV4-20H1 and SUV4-20H2 were generated by CRISPR-Cas9 technology in prostate DU145 cell line. Briefly, DU145 cells were transfected using Jet-PEI with the plasmid pSpCAS9-2A-GFP and the lentiviral vector U6-gRNA2A-TagBFP expressing BFP and guide RNAs targeting SUV4-20H1 (HS5000028803) or SUV4-20H2 (HS5000023102). The guide RNAs were purchased from the Sanger Whole genome CRISPR arrayed library (Merck). Two days after transfection, cells were sorted by flow cytometry for GFP and BFP expression and positive clones were grown for three weeks without selection for Cas9 and RNA guides expression. The cellular clones inactivated for SUV4-20H1 or SUV4-20H2 were then identified by western blotting using antibodies against H4K20 methylation, SUV4-20H1 and SUV4-20H2. The inactivation of SUV4-20H1 or SUV4-20H2 encoding genes was then confirmed by DNA sequencing. The double KO SUV4-20H1 and SUV4-20H2 cells were generated by transfection of SUV4-20H2-KO DU145 cell line with the guide RNA targeting SUV4-20H1.

### Measure of cell proliferation and sensitivity to drug treatments

Cell viability upon drug treatments as indicated was evaluated using resazurin and MTT viability assays and cell proliferation and growth rate were measured with the incucyte cell-by-cell analysis module following the manufacturer recommendations (Sartorius).

### Drug combination screening

For drug combination screening, prostate cancer cell lines were treated with increasing concentration of each drug alone or in combination for three days. The percentage of viable and dead cells were then evaluated using the Celigo flow cytometer after DAPI and propidium iodide (PI) staining and following the manufacture recommendations (Nexcelom). The interaction between the drugs tested *in vitro* was quantified with a concentration matrix test, in which increasing concentration of each single drug were assessed with all possible combinations of the other drugs. For each combination, the percentage of expected growing cells in the case of effect independence was calculated according to the Bliss equation: fuc=fuAfuB, where fuc is the expected fraction of cells unaffected by the drug combination in the case of effect independence, and fuA and fuB are the fractions of cells unaffected by treatment A and B, respectively. The difference between the fuc value and the fraction of living cells in the cytotoxicity test was considered as an estimation of the interaction effect, with positive values indicating synergism and negative values antagonism. The combination index and the efficiency index were calculated based on these values using R software.

### Measure of Synergy and efficiency score and statistical analysis

Cell growth was evaluated using the sulforhodamine B (SRB) assay. Briefly, 1000 cells/well were seeded in 96-well plates. After 24 hours, drugs were added in serial dilution. Cells were incubated for 96 hours, after which, they were fixed in 10% trichloroacetic acid solution (SIGMA), and stained with 0.4 % SRB (SIGMA) solution in 1% acetic acid (FLUKA). The plate was washed three times with 1% acetic acid and fixed SRB was dissolved in 10 mmol/L Tris-Base solution (Trizma®base SIGMA). The absorbance was read at 560 nm using the PHERAstar FS plate reader (BMG LABTECH). The percentage of living cells after incubation with each drug alone or in combination was then calculated. The interaction between the drugs tested in vitro was investigated with a concentration matrix test, in which increasing concentration of each single drug were assessed with all possible combinations of the other drugs (Tosi et al, 2018). For each combination, the percentage of expected growing cells in the case of effect independence was calculated according to the Bliss equation:

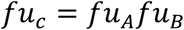

where fu_c_ is the expected fraction of cells unaffected by the drug combination in the case of effect independence, and fu_A_ and fu_B_ are the fractions of cells unaffected by treatment A and B, respectively. The difference between the fu_c_ value and the fraction of living cells in the cytotoxicity test was considered as an estimation of the interaction effect, with positive values indicating synergism and negative values antagonism. Synergy is associated with the color red, additivity with black, and antagonism with green.

According to the approach proposed by Lehar (Lehár et al., 2007, 2008, 2009), a point-by-point calculation of the expected values in case of absence of interaction effect is performed over all the matrix concentration combinations in order to obtain an expected-value matrix. Then, a combination index, CI, is calculated as follows:

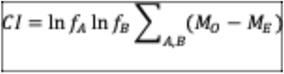

where *M_O_* and *M_E_* are the matrices of the observed and expected values, respectively, and *f_A_* and *f_B_* are the dilution factors for the drugs *A* and *B*. The CI is a positive-gated, effect-weighted volume over the lack of interaction effect (i.e.. Bliss independence), adjusted for variable dilution factors *f_A_* and *f_B_*.

The efficacy Index is calculated as follow:

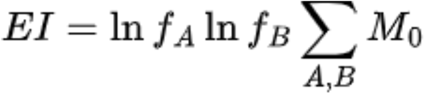

Where M_0_ is the number of dead cells. Combination index represent the level of synergistic effect between two drugs, and the Efficacy Index represent the effect of both drugs on the cell mortality. The two indexes are then plotted in order to visualize the drug combination the most synergistic and efficient.

### RNA sequencing

Total RNA from DU145 cells treated or not with A-196 for 10 days was extracted using the Monarch Total RNA Miniprep Kit (Biolabs), and treated with DNase I treatment to remove genomic DNA contamination. RNA sequencing was performed at MGX Montpellier GenomicX, Montpellier, France. RNA-Sequencing library was prepared using a Stranded mRNA Prep Ligation kit from Illumina. The validation of the libraries was performed by quantifying the concentration and the size of fragments of DNA on Fragment Analyzer (High Sensitivity NGS kit) as well as by qPCR (ROCHE Light Cycler 480). The clustering and sequencing steps were carried out on an Illumina NovaSeq 6000 using the SBS (Sequence by Synthesis) technique using the NovaSeq Reagent Kits. Image analysis and base calling were performed using the NovaSeq Control Software and the Real-Time Analysis component (Illumina). Demultiplexing was performed using Illumina’s conversion software (bcl2fastq, v2.20.0.422). The quality of the raw data was assessed using FastQC (v0.11.9) from the Babraham Institute and the Illumina software SAV (Sequencing Analysis Viewer, v1.1.23). FastQ Screen (v0.15.0) was used to identify potential contamination. The alignment program TopHat2 (v2.1.1), with Bowtie (v2.4.5) ^49^ was used to align the RNA-Seq reads to the human genome (GRCh38, from Ensembl), whose index was built (bowtie2-build) using a set of gene model annotations (gff file downloaded from NCBI on 2022/01/04). For alignment, the TopHat2 was set with the option –library-type fr-firststrand, in accordance to the kit used for library construction, and with options –read-mismatch 6 – read-gap-length –read-edit-dist 6. Otherwise, default TopHat2 options have been used. SAMtools (v1.13) was used to sort and index the alignment files. Then, gene counting was performed with featureCounts (v2.0.3) ^50^. As the data is from a strand-specific assay, the reads have to be mapped to the opposite strand of the gene (‘-s 2’ option). Before statistical analysis, genes with less than 15 reads (cumulating all the analysed samples) were filtered out. Differentially expressed genes were identified using the Bioconductor [4] package DESeq2 (v1.32.0) [5] (R v4.1.1). Data were normalized using the DESeq2 normalization method with a false discovery rate threshold of 0.01 and log FC threshold of 0.69 (50% increase in gene expression). Genes with adjusted p-value below 1% (according to the FDR method from Benjamini-Hochberg) were called differentially expressed (DE). functional enrichment analysis and functional annotation of gene lists were performed with home-made R scripts and DAVID web software from NIH. RNAseq Data have been deposited in the public NIH (SRA) database and available with the accession reference PRJNA1162501. Here the reviewer link: https://dataview.ncbi.nlm.nih.gov/object/PRJNA1162501?reviewer=pt6saodi958qtrd6ni2k06kbnc.

### DNA fiber assays

DNA was labeled for 30 minutes with 25µM 5-iodo-20-deoxyuridine (IdU) (Sigma-Aldrich, 17125) and washed with medium, then treated with 50µM 5-chloro-20-deoxyuridine (CldU) (Sigma-Aldrich, C6891) for 30 minutes. After labelling, cells were collected with trypsin and resuspended in PBS at 500 cells/µL. Cells were lysed with spreading buffer (0.5% SDS, 200 mM Tris-HCl, pH 7.5, and 50 mM EDTA) and DNA fibers were stretched onto glass slides and left to air dry then fixed in methanol/acetic for 10 minutes. After fixation, DNA was denatured with 2.5 M HCl for 1h 30 minutes, and slides were blocked with 6%BSA PBS 0.1% Tween-20 for 1 h. Then, slides were incubated with primary anti-BrdU/IdU (BD Biosciences, 34780) and anti-CldU (Origene, TA190126) for 45 minutes (1/100) followed with secondary antibodies Alexa 488 anti-mouse (Invitrogen, A11017) and anti-rat-Cy3 (Jackson Immuno-research, 712-166153) respectively for 30 minutes (1/100) in PBS 0.1% TritonX-100 at 37°. Slides were mounted with ProLong Diamond antifade (Invitrogen). Fibers were visualized and imaged by Zeiss Axio Imager M2 upright microscope (Zeiss) equipped with an Apochromat 40X objective (NA 1.4, immersion oil). Replication track lengths were analyzed using ImageJ software. Statistical significance was performed using GraphPad Prism by an unpaired t-test. ns: p-value > 0.05; * : p-value < 0.05; ** p-value < 0.01; *** : p-value < 0.001.

### Colony formation assays

Cells were seeded at 50,000 cells/well in 12-well plate. After 24h, cells were treated with combinational concentrations of A-196 and etoposide, Mitoxantrone or Camptothecin for 72h. Following treatment, drugs with medium were removed and cells were harvested and resuspended in fresh medium at 200 cells/ml. Cells were seeded in new 12-well plates and incubated at 37°c for 14 days to allow the formation of colonies. After 14 days, medium was removed, and cells were fixed with methanol for 20 minutes then washed and incubated with crystal violet staining solution for 5 minutes in order to stain the colonies. The number of colonies was counted and statistical significance was evaluated using GraphPad Prism by an unpaired t-test. ns: p-value > 0.05; * : p-value < 0.05; ** p-value < 0.01; *** : p-value < 0.001.

### Protein extraction and immunoblotting

For immunoblot analysis, cells washed with phosphate-buffered saline (PBS) were lysed in 2X SDS Sample Buffer (10 mM Tris-HCl pH6.8, 5% glycerol, 1% SDS, 1 mM DTT, 0.5% β-mercapto-ethanol) at 95°C for 10 minutes. After measuring protein quantity by Bradford, equal amounts of protein were resolved by SDS-PAGE, transferred to a nitrocellulose membrane, and probed with one of the following antibodies: H4K20me1 1/1000 (Cell Signaling Technology, #9724), H4K20me2 1/1000 (Cell Signaling Technology, #9759), H4K20me3 1:1000 (Cell Signaling Technology, #5737), Histone H4 1:1000 (Cell Signaling Technology, #2935), Histone H3 (Cell Signaling Technology, #4499), SUV4-20H1 1:1000, SUV4-20H2 1:1000, Tubulin 1:20000 (Sigma T5168), DNA-PK 1:1000 (Invitrogen, MA513266), phospho-S2056 DNA-PK 1:1000 (Cell Signaling Technology, #6876), ATM 1/1000 (Cell Signaling Technology, #2873), phospho-ATM S1981 1/1000 (Cell Signaling Technology, #13050), ATR 1/:000 (Cell Signaling Technology, #13934), phospho-T1989 ATR 1:1000 (Cell Signaling Technology, #30632), RPA32 1/1000 (Cell Signaling Technology, #35869), phospho-S4/S8 RPA 1:1000 (Cell Signaling Technology, #54762). Secondary antibodies used are the following: Anti-rabbit 1:10000 (Cell Signaling Technology) and anti-mouse 1:10000 (Cell Signaling Technology) coupled to HRP. Protein bands were visualized on X-ray film by electro-chemiluminescence (Immobilon Western HRP Substrate, WBKLS0500, Millipore). The quantification od signal was performed using ImageJ software. Statistical significance was evaluated using GraphPad Prism software by an unpaired t-test. ns: p-value > 0.05; *: p-value < 0.05; ** p-value < 0.01; ***: p-value < 0.001.

### Chromatin fractionation

Cells were resuspended with Buffer A (10 mM Hepes pH 7.9, 10 mM KCl, 1.5 mM MgCl_2_, 0.34 M Sucrose, 10% Glycerol, 1 mM DTT, 0.1% TritonX-100, and protease inhibitors (Roche) and incubated for 2 minutes on ice to extract cytoplasmic soluble proteins. After centrifugation, supernatant containing cytoplasmic protein was collected and cells were washed with the Buffer A without 0.1% TritonX-100. Then, nuclei were resuspended with Buffer B (3 mM EDTA, 0.2 mM EGTA) and incubated for 30 minutes with occasional vortex on ice to extract nuclear soluble proteins. After centrifugation, supernatant containing nuclear soluble proteins was collected and cells were washed with the Buffer B. Then, chromatin was dissolved in 2X SDS Sample Buffer (10 mM Tris-HCl pH6.8, 5% glycerol, 1% SDS, 1 mM DTT, 0.5% β-mercaptoethanol) and resolved by SDS-PAGE before immunoblotting.

### Immunofluorescence and Microscopy analysis

Cells were seeded and treated after 24h with A-196 4µM for 72h. Then, cells were transferred on coverslips with maintenance of A-196 and grown to 50% confluency to be treated with etoposide for 24h. Following this treatment, cells were washed with medium and a fresh medium was added with or without the maintenance of A-196. At the indicated time, cells were pre-extracted with ice-cold PBS containing 0.1% Triton-X 100 for 1-2 minutes and fixed with 4% paraformaldehyde (Thermo Scientific) diluted in PBS for 15 minutes at room temperature (RT) after 0, 6, 24, and 48hr of etoposide removal. After fixation, cells were permeabilized with a solution of PBS with 0.25% Triton-X 100 for 10 minutes at RT. Then, cells were incubated for 1h at RT in PBS solution complemented with 5% BSA and 0.1% Tween20 prior to incubation overnight at 4°C with the following primary antibodies : γ-H2AX 1:500 (Millipore, JBW301), 53BP1 1:500 (Cell Signaling Technology, #4937), RPA70 1:100 (Cell Signaling, #2267), RAD51 1:500 (Millipore, PC130). On the following day, cells were washed with PBS solution containing 0.5% Tween-20 and incubated with appropriate secondary mouse and rabbit antibodies coupled to Alexa Fluor 568 or 488 (Thermo Fisher Scientific) diluted (1:1000) in PBS solution with 5% BSA and 0.1% Tween-20 for 2 hours at RT. DNA was counterstained with 0.1 µg/ml of DAPI (DNA intercalator, 4’,6-Diamidino-2-Phenylindole, D1306, Invitrogen). Coverslips were mounted with ProLong Diamond Antifade (Invitrogen). Images were acquired with an ORCA-Flash4.0 LT+ Digital CMOS camera (C1140-42U30, Hamamatsu) controlled by Zen acquisition software (Zeiss) using Zeiss Axio Imager M2 upright microscope (Zeiss) equipped with an Apochromat 63X objective (NA 1.4, immersion oil). Images were prepared using Zen software and nuclear foci were quantified using CellProfiler^TM^ software.

Representative images were prepared using Photoshop software (Adobe). Statistical significance was evaluated using GraphPad Prism by an unpaired t-test or two-tailed Student t-test. ns: p-value > 0.05; * p-value < 0.05; ** p-value < 0.01; *** p-value < 0.001.

### Flow cytometry analysis

Prior fixation, cells were incubated with BrdU (300µM, Sigma-Aldrich) for 1 hour. Cells were then fixed with an ice-cold 70% ethanol solution and permeabilized with 0,2% Triton X-100 for 10min. DNA was then denatured with 0,2N HCl before incubation with BrdU antibody (1:200, BD, Biosciences) diluted in PBS solution with 0,2% Tween20 and 1% BSA for 1 hour at RT followed by 1 hour incubation with an FITC-conjugated antibody (BD, Biosciences; 1;300). DNA was then counterstained by 12 hours incubation with 2 μg/ml 7-Amino-Actinomycin D (7AAD, Sigma-Aldrich) in presence of RNAse (100 μg/ml). FACS acquisition were carried out with Gallios flow cytometer (Beckman Coulter). FACS data were analyzed using FlowJo software (BD Biosciences). Statistical significance was evaluated using GraphPad Prism software by an unpaired t-test. ns: p-value > 0.05; *: p-value < 0.05; ** p-value < 0.01; *** p-value < 0.001.

### Pulse Field Gel Electrophoresis

Cells were seeded and treated after 24h with A-196 4µM for 72h. Then, cells were transferred in new plates with maintenance of A-196 and grown to 50% confluency to be treated with etoposide for 3 hours. After treatment, cells were collected with trypsinization and washed once with PBS. 10^6^ cells were then melted in 0.5% agarose insert. After solidification, these inserts were incubated in lysis buffer (100 mM EDTA pH8, 0.2% sodium deoxycholate, 1% sodium lauryl sarcosine, 1 mg/ml Proteinase K) at 37 °C for 48h and then washed 2-4 times with the following buffer: 20 mM Tris pH 8, 50 mM EDTA pH 8. Before loading onto a 0.9% agarose prepared in 0.25X TBE, DNA is separated by pulsed-field gel electrophoresis for 24 hours (Biometra Rotaphor 8 System, 23h; interval: 30-5 s log; angle: 120-110 linear; voltage: 180-120 V log, 13°C). The gel was subsequently stained with ethidium bromide for analysis. The quantification was performed using ImageLab software. Statistical significance was evaluated using GraphPad Prism software by an unpaired t-test. ns: p-value > 0.05; * p-value < 0.05; ** p-value < 0.01; *** p-value < 0.001.

### Heparin-based extraction for TOP2cc immunodetection

Cells were plated and treated after 24h with A-196 4µM for 72h. Then, cells were transferred into a 6-well plate with maintenance of A-196 and grown to 50% confluency to be treated with etoposide for 3h. For immunoblotting analysis, cells were harvested with trypsin, and washed with cold PBS. After gentle centrifugation, pellets were resuspended in lysis buffer (150 mM NaCl, 1 mM EDTA, 0.5% IGEPAL CA-630, 2X HALT Protease and Phosphatase Inhibitor Cocktail [Thermo Fisher Scientific] 20 mM Tris-HCl, pH 8.0) complemented with 100 U/mL Heparin (H3393, Sigma-Aldrich) for 15 minutes on ice. After the incubation, lysates were centrifuged at 15000 rpm for 5 minutes at 4°C and pellets were resuspended again with lysis buffer. To facilitate the migration of the protein in immunoblotting, two cycles of sonication were performed to degrade the DNA present in the extracts using Bioruptor Pico (Diagenode) (2 cycles – 30 sec ON and 30 sec Off each). Protein concentration was then determined with Nanodrop by measuring the absorbance at 280 nm. Before loading, heparin-based extracts were diluted with 2X SDS Sample Buffer (10 mM Tris-HCl pH6.8, 5% glycerol, 1% SDS, 1 mM DTT, 0.5% β-mercaptoethanol) and Immunoblotting was performed with the following antibodies: anti-Topoisomerase IIα 1:1000 (Santa Cruz sc-365916) and anti-histone H3 1:2500 (Cell signaling Technology #4499).

### In vivo Animal studies

The Institute of Animal Care and French APAFIS committee approved all mouse protocols used in this study (French National Agreement #28412). One million of DU145 cells and SUV4-20Hs knock-out cells were subcutaneously injected into intact athymic nude male mice. Treatment was initiated when tumor size reached ∼100 mm3 via intraperitoneal injection (i.p) two times per week. For the combination therapy experiment with DU145 xenografts, mice were randomized into four treatment groups: Vehicle (20% DMSO/corn oil), A-196 (7.5 mg/kg), etoposide (2.5 mg/kg), and combination of the two drugs. For the experiment with wild-type and SUV4-20Hs knock-out DU145 xenografts, male mice were randomized into different treatment groups and treated with DMSO or with 2.5 mg/kg or 6mg/kg of etoposide every three days. Mice were weighed weekly to monitor for toxicity and tumor growth was assessed by serial caliper measurements every three days. Statistical significance was evaluated using GraphPad Prism software by an unpaired t-test. ns: p-value > 0.05; *: p-value < 0.05; ** p-value < 0.01; ***: p-value < 0.001.

## Supporting information

table 1 RNAseq

table 2 RNAseq

## ACKNOWLEDGMENTS

This work was supported by several grants from SIRIC Montpellier Cancer (INCa-DGOS-INSERM-ITMO Cancer-18004), Ligue Contre le Cancer Languedoc-Rousillon, MUSE Innovation program of Montpellier University (project CAPRE) and Institut National du Cancer (INCA PLBIO24-148). MGX acknowledges financial support from France Génomique National infrastructure and Agence Nationale pour la Recherche (ANR-10-INBS-09). Institutional Support was provided by the Institut National de la Santé et de la Recherche Médicale (INSERM), Montpellier University and by the Centre National de la Recherche Scientifique (CNRS). F.A. was supported by a PhD fellowship provided by Montpellier University, CNRS-Liban and by the Ligue Nationale Contre le Cancer (LNCC).

## AUTHORS CONTRIBUTION

EJ designed the project, VB, RAM and EJ supervised the research. FA and JP performed the in vitro experiments. FA, MT, JP and AT performed the in vivo experiments. EJ, SG and XM performed RNA sequencing. FA, VB, RAM and EJ analyzed the data. CR, CG, FC, MD and DT provided conceptual advices. FA, VB and EJ wrote the paper. FA, VB, CR, PP, CG, RAM and EJ contributed to manuscript editing.

## LEGENDS OF SUPPLEMENTARY FIGURES

**Figure S1: Clinical features associated with SUV4-20H2 expression in prostate tumors. (A)** Bar plot representing the mRNA expression of SUV4-20H2 according to the Gleason Score (GS). Error bars represent the minimum and maximum level of expression, ***P < 0.0001. (**B**) Dot plot presenting SUV4-20H2 mRNA expression in two age categories of prostate cancer patients. Patients from 41-60 years (red dots) and patients from 61-80 years (orange dots). Mean values (black lines). **, P < 0.001. (**C**) Dot plot representing SUV4-20H2 mRNA expression in prostate cancer patients with TMPRSS2 structural variants and (**D**) with SPOP mutations. No mutation (red dots) and with mutation (orange dots). Mean values (black lines). ns: non-significant. (**E**) Dot plot representing SUV4-20H2 mRNA expression in prostate cancer patients with or without TP53 mutations. **, P < 0.001 (**F**) Dot plot representing SUV4-20H2 mRNA expression in two stages of Neoplasm Disease Lymph Node. N0 (red dots) and N1 (orange dots). Mean values (black lines). *, P < 0.01. (**G**) Dot plot representing SUV4-20H2 mRNA expression in prostate cancer patients with or without ERG structural variants. No structural variant (red dots) and with structural variants (orange dots). Mean values (black lines), *, P < 0.01.

**Figure S2: SUV4-20H2 upregulation correlates with genetic instability in the cohort of prostate tumors in TCGA database.** Dot plot showing the fraction Genome Altered (top left), the MSI MANTIS Score (top middle), the Aneuploidy Score (top right), the Buffa Hypoxia Score (bottom left), and the Winter Hypoxia Score (bottom right) in patients with low SUV4-20H2 expression (red dots) and high SUV4-20H2 expression (blue dots) levels from the prostate TCGA database. Mean values (black lines). **, P < 0.001; ***, P < 0.0001.

**Figure S3: Loss of SUV4-20H1 or SUV4-20H2 accelerates replication fork rates.** Dot plot represents the IdU (green) and CldU (red)-labeled replication track lengths in DU145 cells inactivated for SUV4-20H1 or SUV4-20H2 using Crispr-Cas9 approach, n=3, 100 replication tracks per condition, p-value as indicated in the graph.

**Figure S4: Features of the lethal synergy between the SUV4-20Hs inhibitor A-196 and etoposide in metastatic prostate cancer cell lines. (A)** Scatter plot representing the combination index (y) and the efficiency index (x) of combination treatment with increasing concentrations of A-196 and etoposide for 72h in 22RV1, DU145, PC3, and LNCaP cell lines. Each dot represents one experiment. **(B)** Representative viability matrix (top) and synergy scores (bottom) for DU145 and PC3 cell lines after 72h of treatment with increasing doses of A-196 and etoposide or with increasing doses of EZH2i (GSK126) and etoposide as indicated. Synergy occurs when score values are greater than 20 (Red), additivity occurs when score values are between −20 and 20 (Black), and antagonism occurs when score values are less than −20 (Green). Score and color intensity are dependent on the degree of synergy, antagonism or additivity.

**Figure S5: The loss of SUV4-20H proteins increases DNA damage in response to etoposide similarly to their inhibition with A-196 compound.** Upper panel, Representative images of 53BP1 and γ-H2AX staining in DU145 WT and DU145 double KOSUV4-20H clones treated with 0.45μM of etoposide or not (DMSO) for 24h as indicated. Scale bar = 10μm. Lower panel, Violin plot representing the number of 53BP1 (Left) and γ-H2AX (right) foci per nucleus of DU145 WT and DU145 double KOSUV4-20H clones treated as indicated. Median (plain line), quartile (dashed line). n = 1, >200 cells per condition, *p-value < 0.01, **p-value <0.001, ***p-value< 0.0001.

**Figure S6: RPA and DNAPKcs remain phosphorylated in A196-treated cells after etoposide removal.** DU145 cell line was treated with 4μM A-196 or not (DMSO) for 72h Immunoblot analysis showing the maintenance of RPA and DNAPKcs phosphorylation 6, 24, and 48h after 24 hours exposure with 0.45μM of etoposide in DU145 cell line that was previously treated with 4μM A-196 or not (DMSO) uding 72 hours in order to induce H4K20me reprogramming.

**Figure S7: The pharmacological inhibition of SUV4-20Hs enhances etoposide responses in castration-resistant 22RV1 prostate cell lines. (A)** Representative images of γ-H2AX staining in 22RV1 cell line treated with A-196 4μM or not (DMSO) for 72h, then etoposide 0.63μM was added for an additional 24h as indicated. Scale bar = 10μm. **(B)** Immunoblot analysis showing the expression levels of indicated proteins in 22RV1 cell line treated with A-196 4μM or not (DMSO) for 72h, etoposide 0.63μM was added for additional 24h when indicated (+). Total DNA-PKcs, ATM, and ATR are used as a loading control, n=2. **(C)** Bar plot representing the number of γ-H2AX foci per nucleus of 22RV1 cell line treated with A-196 4μM or not (DMSO) for 72h, followed by etoposide 0.63μM addition for 24h (0h of etoposide removal) and after 6, 24, and 48h of etoposide removal as indicated. A-196 is maintained when indicated (+), n=2, >200 cells per condition.

**Figure S8: A 24-hour treatment with a harmless dose of etoposide induces S/G2 arrest and cell death in A-196-treated DU145 cells. (A)** Cell cycle distribution of DU145 cells 6, 24, and 48h after 24 hours treatment with etoposide (0.45μM) and treatment or not with A-196 (4μM) 72 hours before etoposide incubation. Total DNA was stained with 7-AAD, and nascent DNA was labeled with BrdU, n=3. **(B)** Bar plot representing the % of sub-G1 cells for DU145 cell line treated or not as indicated, n=3, ns: non-significant, * p-value< 0.01, ** p-value < 0.001, ***p-value < 0.0001.

