## supplementary figures for "Targeting SUV4-20H epigenetic enzymes as therapeutic strategy for enhancing topoisomerase II poisoning in prostate cancer"

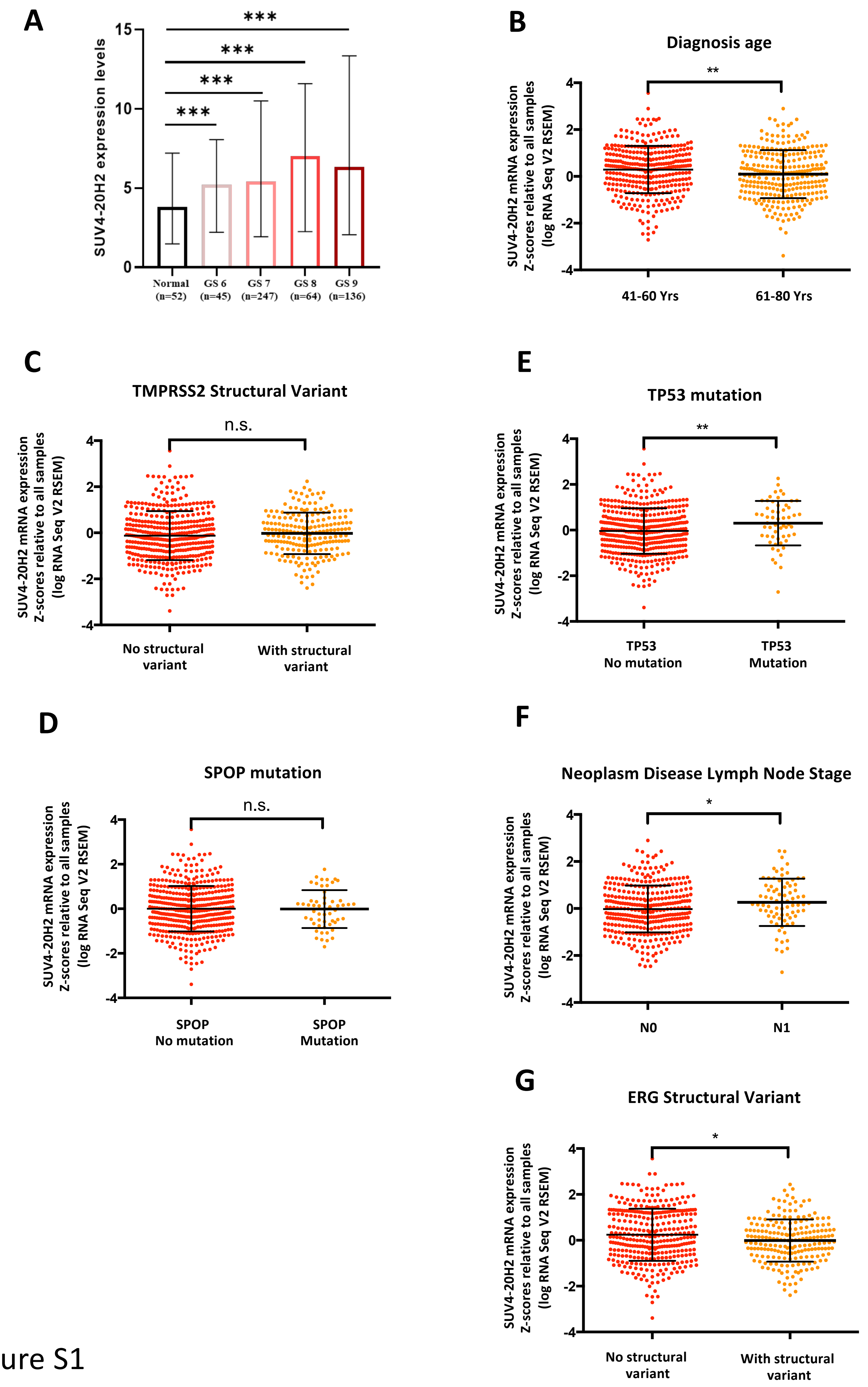


Figure S1

Alhourani F et al.

- **Figure S1: Clinical features associated with SUV4-20H2 expression in prostate tumors. (A)** Bar plot representing the mRNA expression of SUV4-20H2 according to the Gleason Score (GS). Error bars represent the minimum and maximum level of expression, ***P < 0.0001. (**B**) Dot plot presenting SUV4-20H2 mRNA expression in two age categories of prostate cancer patients. Patients from 41-60 years (red dots) and patients from 61-80 years (orange dots). Mean values (black lines). **, P < 0.001. (**C**) Dot plot representing SUV4-20H2 mRNA expression in prostate cancer patients with TMPRSS2 structural variants and (**D**) with SPOP mutations. No mutation (red dots) and with mutation (orange dots). Mean values (black lines). ns: non-significant. (**E**) Dot plot representing SUV4-20H2 mRNA expression in prostate cancer patients with or without TP53 mutations. **, P < 0.001 (**F**) Dot plot representing SUV4-20H2 mRNA expression in two stages of Neoplasm Disease Lymph Node. N0 (red dots) and N1 (orange dots). Mean values (black lines). *, P < 0.01. (**G**) Dot plot representing SUV4-20H2 mRNA expression in prostate cancer patients with or without ERG structural variants. No structural variant (red dots) and with structural variants (orange dots). Mean values (black lines), *, P < 0.01.

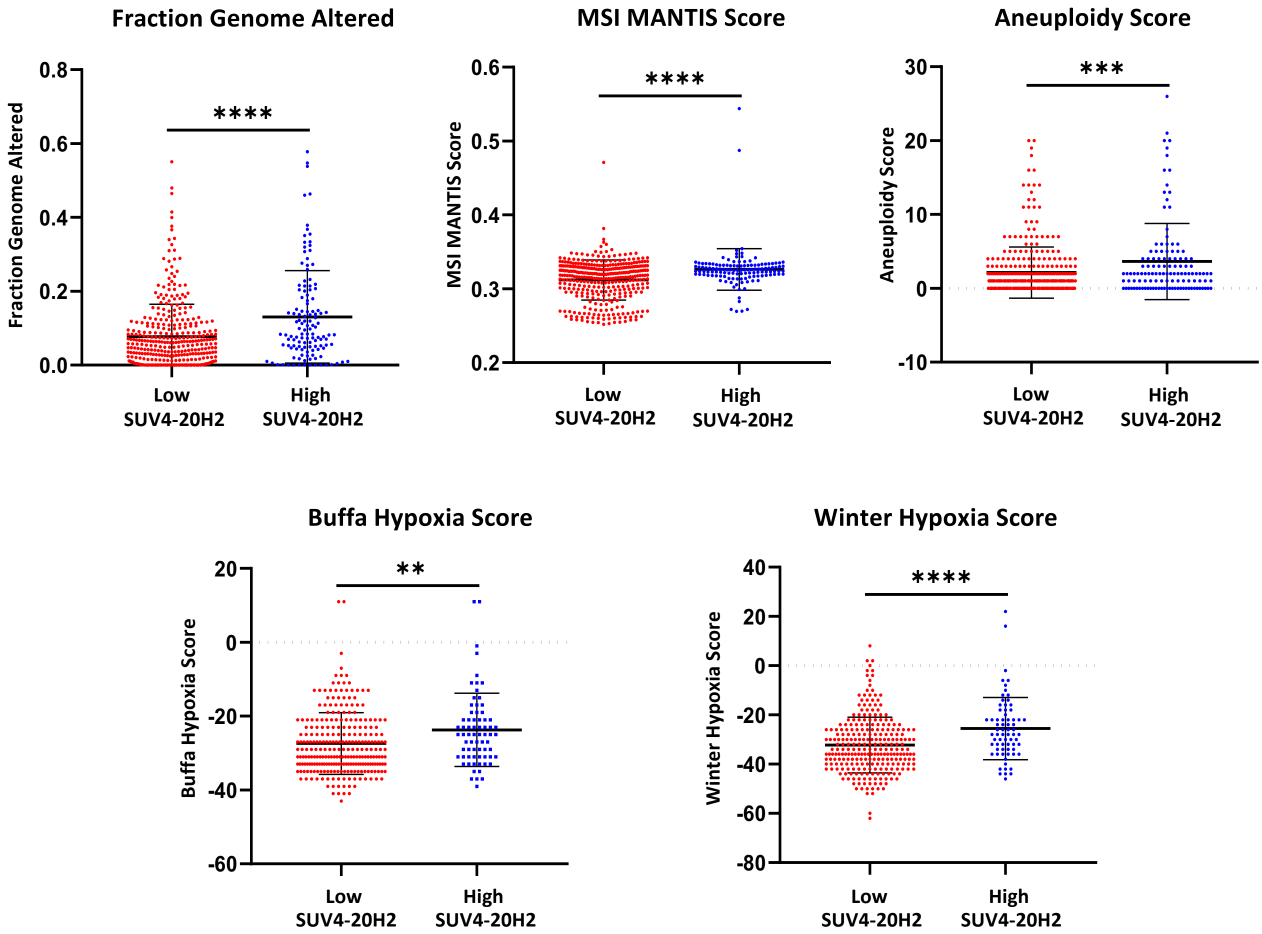


**Figure S2: SUV4-20H2 upregulation correlates with genetic instability in the cohort of prostate tumors in TCGA database.** Dot plot showing the fraction Genome Altered (top left), the MSI MANTIS Score (top middle), the Aneuploidy Score (top right), the Buffa Hypoxia Score (bottom left), and the Winter Hypoxia Score (bottom right) in patients with low SUV4-20H2 expression (red dots) and high SUV4-20H2 expression (blue dots) levels from the prostate TCGA database. Mean values (black lines). **, P < 0.001; ***, P < 0.0001.

Figure S2

Alhourani F et al.

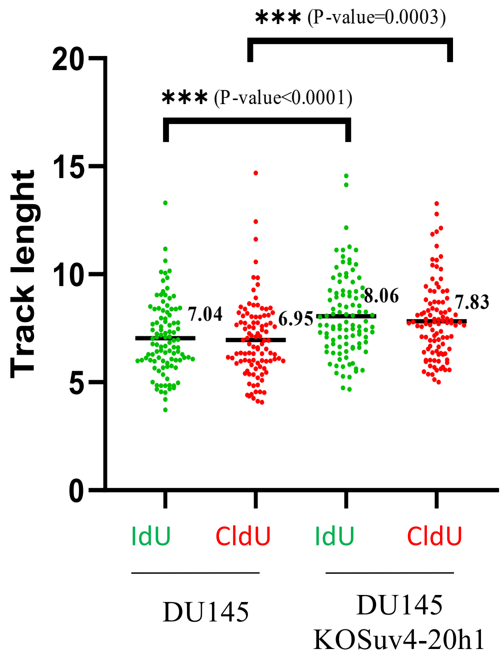


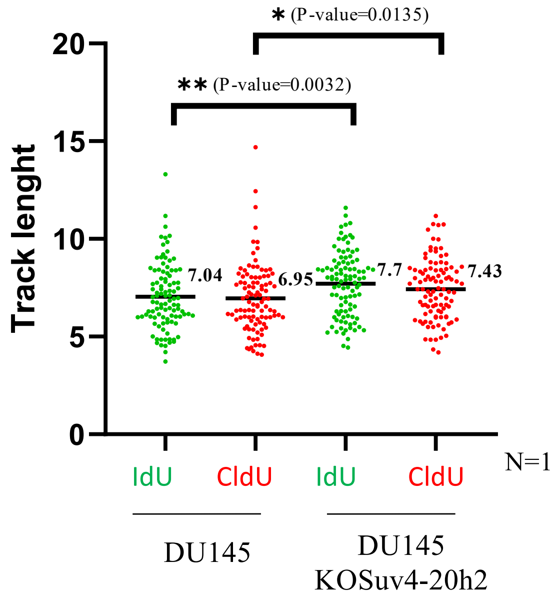


**Figure S3: Loss of SUV4-20H1 or SUV4-20H2 accelerates replication fork rates.** Dot plot represents the IdU (green) and CldU (red)-labeled replication track lengths in DU145 cells inactivated for SUV4-20H1 or SUV4-20H2 using Crispr-Cas9 approach, n=3, 100 replication tracks per condition, p-value as indicated in the graph.

Figure S3

Alhourani F et al.

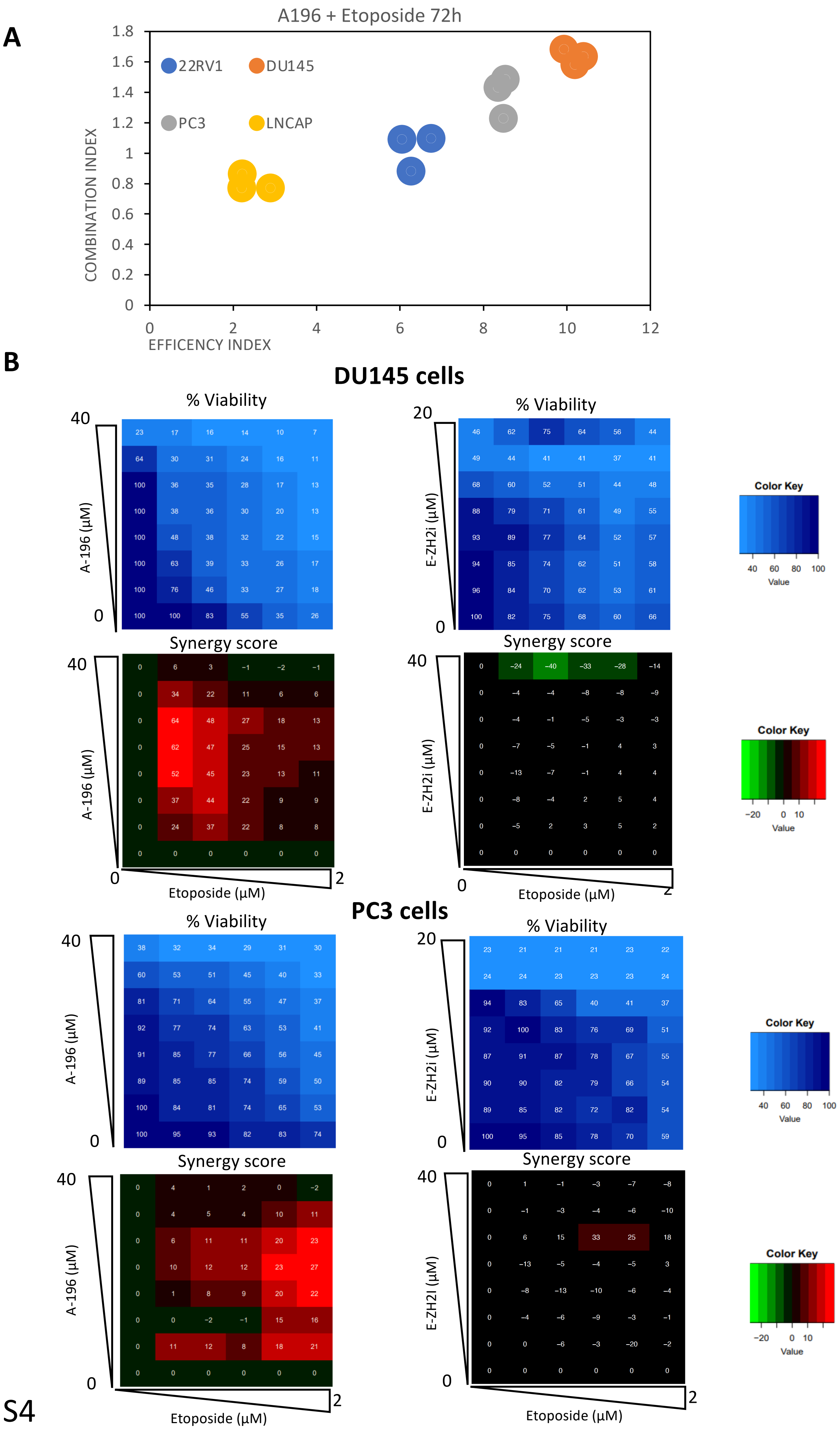


Figure S4

Alhourani F et al.

**Figure S4 : Features of the lethal synergy between the SUV4-20Hs inhibitor A-196 and etoposide in metastatic prostate cancer cell lines. (A)** Scatter plot representing the combination index (y) and the efficiency index (x) of combination treatment with increasing concentrations of A-196 and etoposide for 72h in 22RV1, DU145, PC3, and LNCaP cell lines. Each dot represents one experiment. **(B)** Representative viability matrix (top) and synergy scores (bottom) for DU145 and PC3 cell lines after 72h of treatment with increasing doses of A-196 and etoposide or with increasing doses of EZH2i (GSK126) and etoposide as indicated. Synergy occurs when score values are greater than 20 (Red), additivity occurs when score values are between -20 and 20 (Black), and antagonism occurs when score values are less than -20 (Green). Score and color intensity are dependent on the degree of synergy, antagonism or additivity.

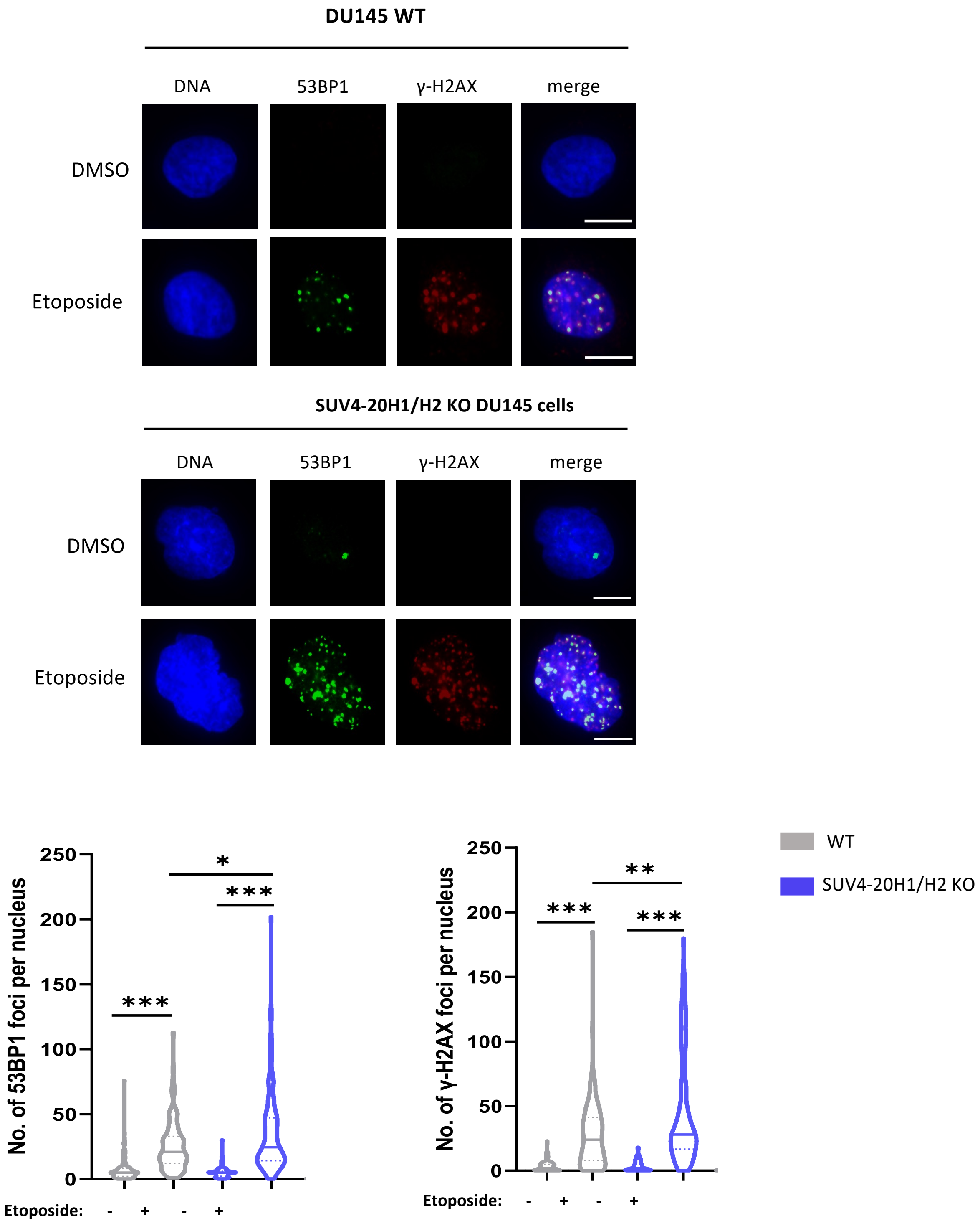


**Figure S5: The loss of SUV4-20H proteins increases DNA damage in response to etoposide similarly to their inhibition with A-196 compound.** Upper panel, Representative images of 53BP1 and γ-H2AX staining in DU145 WT and DU145 double KOSUV4-20H clones treated with 0.45µM of etoposide or not (DMSO) for 24h as indicated. Scale bar = 10µm. Lower panel, Violin plot representing the number of 53BP1 (Left) and γ-H2AX (right) foci per nucleus of DU145 WT and DU145 double KOSUV4-20H clones treated as indicated. Median (plain line), quartile (dashed line). n = 1, >200 cells per condition, *p-value < 0.01, **p-value <0.001, ***p-value< 0.0001.

### Figure S5

Alhourani F et al.

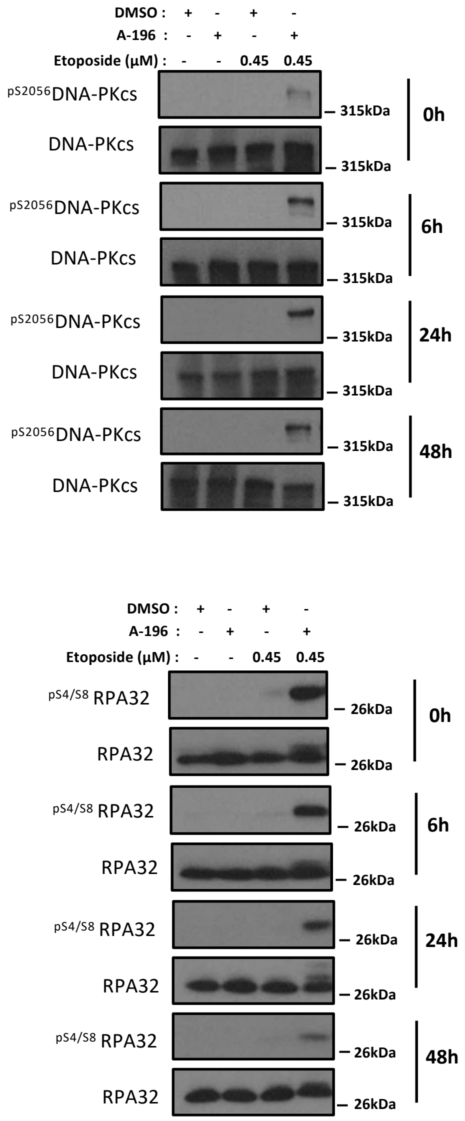


**Figure S6: RPA and DNAPKcs remain phosphorylated in A196-treated cells after etoposide removal.** DU145 cell line was treated with 4µM A-196 or not (DMSO) for 72h Immunoblot analysis showing the maintenance of RPA and DNAPKcs phosphorylation 6, 24, and 48h after 24 hours exposure with 0.45µM of etoposide in DU145 cell line that was previously treated with 4µM A-196 or not (DMSO) uding 72 hours in order to induce H4K20me reprogramming.

Figure S6

Alhourani F et al.

A

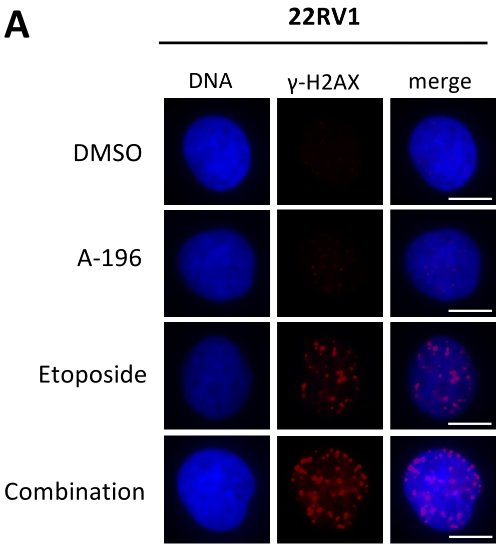


B

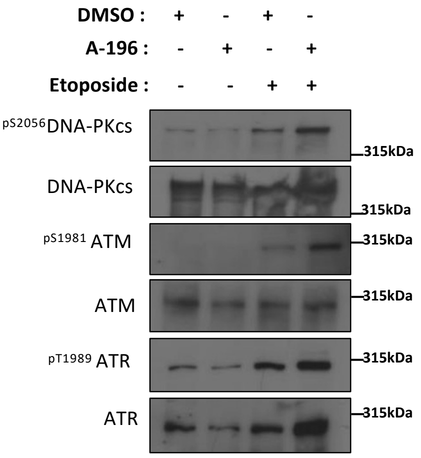


C

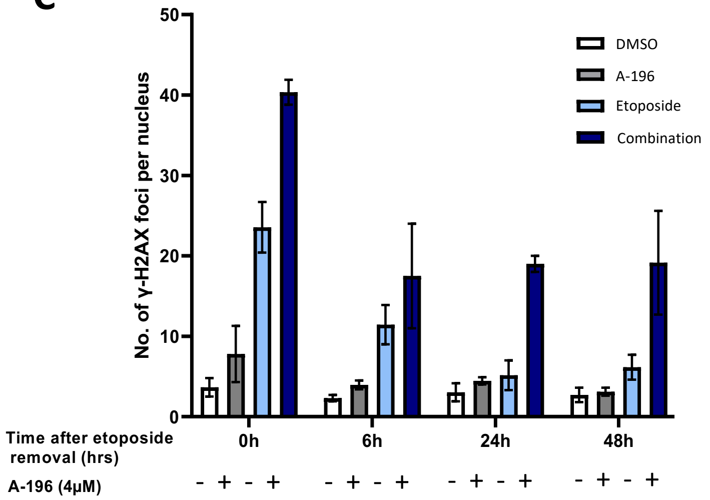


### Figure S7

Alhourani F et al.

**Figure S7: The pharmacological inhibition of SUV4-20Hs enhances etoposide responses in castration-resistant 22RV1 prostate cell lines. (A)** Representative images of γ-H2AX staining in 22RV1 cell line treated with A-196 4μM or not (DMSO) for 72h, then etoposide 0.63μM was added for an additional 24h as indicated. Scale bar = 10μm. **(B)** Immunoblot analysis showing the expression levels of indicated proteins in 22RV1 cell line treated with A-196 4μM or not (DMSO) for 72h, etoposide 0.63μM was added for additional 24h when indicated (+). Total DNA-PKcs, ATM, and ATR are used as a loading control, n=2. **(C)** Bar plot representing the number of γ-H2AX foci per nucleus of 22RV1 cell line treated with A-196 4μM or not (DMSO) for 72h, followed by etoposide 0.63μM addition for 24h (0h of etoposide removal) and after 6, 24, and 48h of etoposide removal as indicated. A-196 is maintained when indicated (+), n=2, >200 cells per condition.

**A**

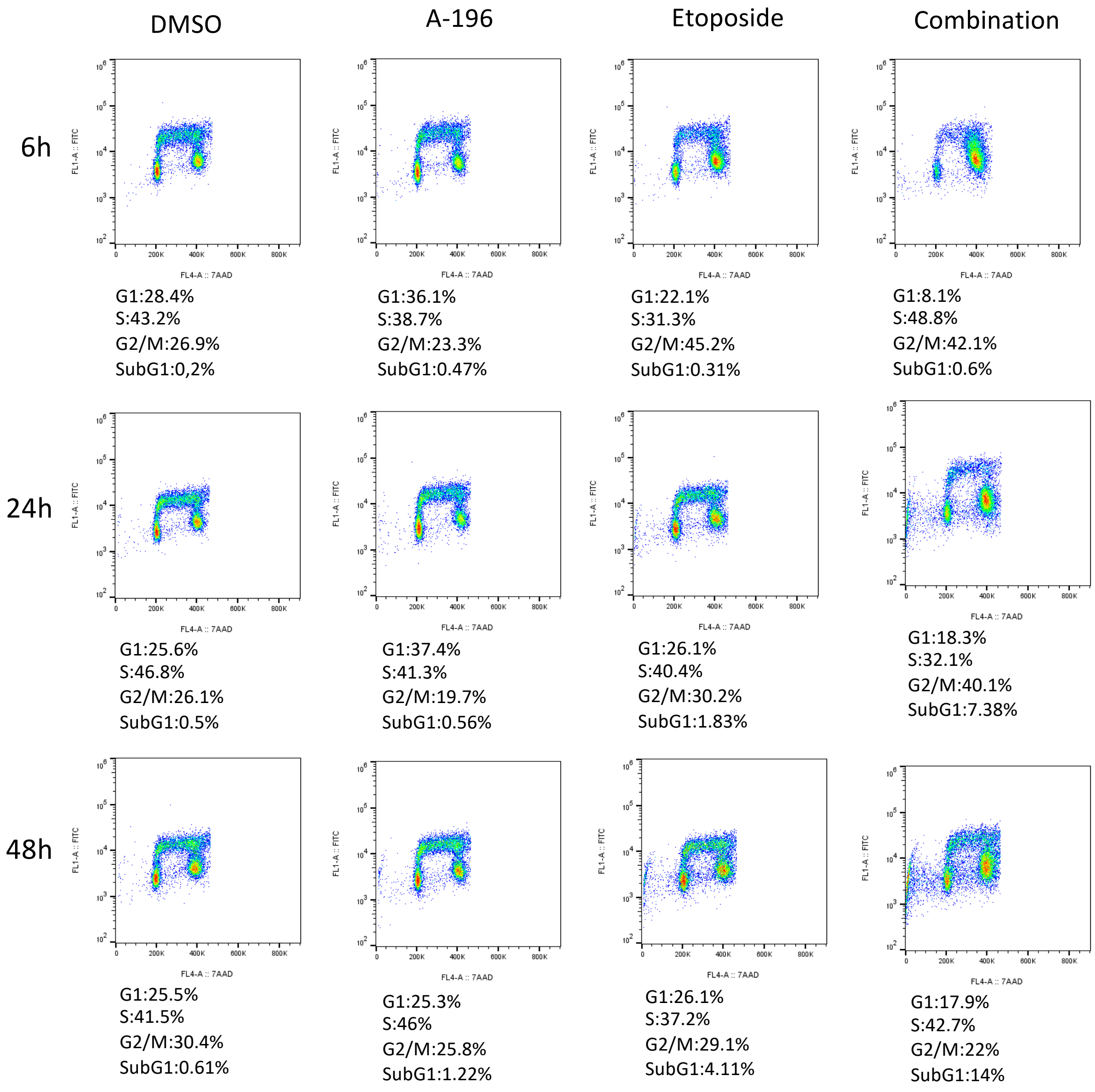


**B**

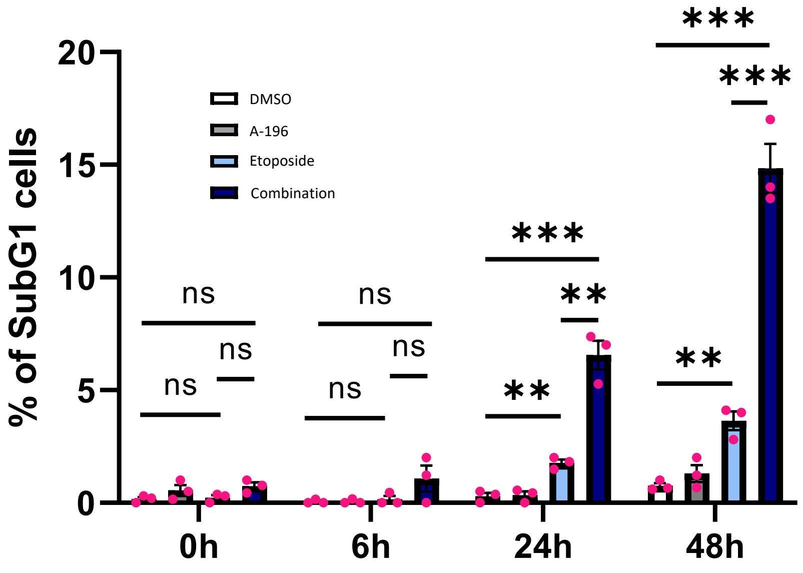


Figure S8

Alhourani F et al.

**Figure S8: A 24-hour treatment with a harmless dose of etoposide induces S/G2 arrest and cell death in A-196-treated DU145 cells. (A)** Cell cycle distribution of DU145 cells 6, 24, and 48h after 24 hours treatment with etoposide (0.45µM) and treatment or not with A-196 (4µM) 72 hours before etoposide incubation. Total DNA was stained with 7-AAD, and nascent DNA was labeled with BrdU, n=3. **(B)** Bar plot representing the % of sub-G1 cells for DU145 cell line treated or not as indicated, n=3, ns: non-significant, * p-value< 0.01, ** p-value < 0.001, ***p-value < 0.0001.
