## Supplementary material for "Targeting SUV4-20H epigenetic enzymes as therapeutic strategy for enhancing topoisomerase II poisoning in prostate cancer": table 1 RNAseq

| feature | GENE_ID | baseMean | log2 (FC) | lfcSE | stat | pvalue | padj |
|---|---|---|---|---|---|---|---|
| LOC106699570 | 106699570 | 17.42473228 | -1.904069604 | 0.50286723 | 3.7864261 | 0.00015283 | 0.00216664 |
| FOS | 2353 | 183.8014951 | -1.342251894 | 0.189906621 | 7.06795734 | 1.57E-12 | 3.25E-10 |
| DMBT1 | 1755 | 45.46205698 | -1.263428103 | 0.330732387 | 3.82009187 | 0.0001334 | 0.00193305 |
| ADCY10P1 | 221442 | 55.10428121 | -1.174711118 | 0.272116831 | 4.31693664 | 1.58E-05 | 0.00035114 |
| NPNT | 255743 | 315.0236295 | -1.067635284 | 0.126946054 | 8.41014946 | 4.09E-17 | 2.47E-14 |
| SLC6A3 | 6531 | 400.2994783 | -1.002281214 | 0.135050473 | 7.42153056 | 1.16E-13 | 3.10E-11 |
| CDRT1 | 374286 | 89.25968942 | -0.88722126 | 0.208786431 | 4.24942012 | 2.14E-05 | 0.00045037 |
| PIK3R3 | 8503 | 122.177714 | -0.87831471 | 0.179999623 | 4.87953638 | 1.06E-06 | 3.82E-05 |
| DOK3 | 79930 | 89.73894095 | -0.862781595 | 0.215101644 | 4.01104138 | 6.05E-05 | 0.00103406 |
| SLC43A2 | 124935 | 1445.525001 | -0.850219224 | 0.116646136 | 7.28887605 | 3.13E-13 | 7.60E-11 |
| KLF9 | 687 | 561.014703 | -0.831657626 | 0.11168078 | 7.44673904 | 9.57E-14 | 2.67E-11 |
| SPTBN5 | 51332 | 155.2348596 | -0.819106354 | 0.173870577 | 4.71101188 | 2.46E-06 | 7.64E-05 |
| TRIM54 | 57159 | 79.47157666 | -0.799459098 | 0.228800467 | 3.49413228 | 0.00047561 | 0.00520885 |
| DOCK11 | 139818 | 2305.752769 | -0.798677069 | 0.090460799 | 8.82898533 | 1.06E-18 | 8.84E-16 |
| MAN1A1 | 4121 | 575.9081358 | -0.795760118 | 0.115407457 | 6.8952227 | 5.38E-12 | 9.32E-10 |
| KCNH1-IT1 | 100874296 | 81.42937364 | -0.789676957 | 0.226177253 | 3.4914075 | 0.00048048 | 0.00525463 |
| MEGF9 | 1955 | 9276.57 | -0.751134724 | 0.094104477 | 7.98192333 | 1.44E-15 | 5.83E-13 |
| TP53INP1 | 94241 | 1089.326995 | -0.73598954 | 0.107057073 | 6.87473996 | 6.21E-12 | 1.03E-09 |
| BMF | 90427 | 585.1229806 | -0.732894095 | 0.133468401 | 5.49114315 | 3.99E-08 | 2.41E-06 |
| PRUNE2 | 158471 | 221.8443018 | -0.725537499 | 0.183014094 | 3.96438047 | 7.36E-05 | 0.00121463 |
| SLC17A8 | 246213 | 132.0300483 | -0.721163324 | 0.171828611 | 4.1969921 | 2.70E-05 | 0.00054787 |
| TAF9B | 51616 | 298.0540008 | -0.71872771 | 0.137573548 | 5.22431615 | 1.75E-07 | 8.44E-06 |
| PER3 | 8863 | 188.2085307 | -0.71398397 | 0.156438552 | 4.5639899 | 5.02E-06 | 0.00013678 |
| ABCC4 | 10257 | 3193.606927 | -0.700513497 | 0.080049118 | 8.75104583 | 2.11E-18 | 1.68E-15 |
| TNFRSF21 | 27242 | 2824.396757 | -0.699777502 | 0.082491467 | 8.4830289 | 2.19E-17 | 1.38E-14 |
| ATP11C | 286410 | 3190.79314 | -0.695962027 | 0.08716572 | 7.98435474 | 1.41E-15 | 5.83E-13 |
