## Supplementary material for "Targeting SUV4-20H epigenetic enzymes as therapeutic strategy for enhancing topoisomerase II poisoning in prostate cancer": table 2 RNAseq

| feature | gene_ID | baseMean | Log2(FC) | lfcSE | stat | pvalue | pADJ |
|---|---|---|---|---|---|---|---|
| ROR2 | 4920 | 18.45223854 | 2.623863042 | 0.52173683 | -5.029093 | 4.93E-07 | 2.04E-05 |
| GPR1 | 2825 | 11.24881559 | 2.552345225 | 0.65141651 | -3.9181463 | 8.92E-05 | 0.001405156 |
| SRGN | 5552 | 16.62868286 | 2.33328246 | 0.52668872 | -4.4300977 | 9.42E-06 | 0.000229313 |
| MIG7 | 9878 | 63.5159673 | 2.046565684 | 0.26404106 | -7.7509372 | 9.12E-15 | 2.92E-12 |
| LINGO1 | 84894 | 24.12753368 | 1.946457058 | 0.4646131 | -4.1894149 | 2.80E-05 | 0.000564973 |
| PRSS8 | 5652 | 25.59028959 | 1.917053474 | 0.41207674 | -4.6521759 | 3.28E-06 | 9.57E-05 |
| KRT81 | 3887 | 648.5026397 | 1.896571464 | 0.12574093 | -15.083167 | 2.09E-51 | 1.57E-47 |
| PSG5 | 5673 | 34.90040043 | 1.849731777 | 0.35076787 | -5.2733786 | 1.34E-07 | 6.80E-06 |
| SLC2A3 | 6515 | 40.86456767 | 1.800377519 | 0.3148504 | -5.7181998 | 1.08E-08 | 7.41E-07 |
| TNS4 | 84951 | 106.7196892 | 1.788654414 | 0.21641945 | -8.2647582 | 1.40E-16 | 7.27E-14 |
| IGFBP3 | 3486 | 1442.63088 | 1.72902404 | 0.0831751 | -20.78776 | 5.59E-96 | 8.42E-48 |
| ALPG | 251 | 154.5802133 | 1.725943396 | 0.2118319 | -8.1477031 | 3.71E-16 | 1.80E-13 |
| PEAR1 | 375033 | 61.01473851 | 1.720203329 | 0.26602441 | -6.4663365 | 1.00E-10 | 1.23E-08 |
| COL1A2 | 1278 | 37.48349737 | 1.690507785 | 0.32982156 | -5.1255224 | 2.97E-07 | 1.30E-05 |
| PRRT4 | 401399 | 82.69868077 | 1.685179781 | 0.22712222 | -7.4197047 | 1.17E-13 | 3.10E-11 |
| KREMEN2 | 79412 | 100.5774223 | 1.64405213 | 0.219774 | -7.4806489 | 7.40E-14 | 2.14E-11 |
| GKN2 | 200504 | 325.9291107 | 1.638989896 | 0.13386133 | -12.243938 | 1.81E-34 | 6.82E-31 |
| SCUBE2 | 57758 | 19.16037968 | 1.62410663 | 0.45637362 | -3.5587215 | 0.00037266 | 0.004303488 |
| TNFSF15 | 9966 | 34.82894534 | 1.617358718 | 0.34263226 | -4.7203925 | 2.35E-06 | 7.34E-05 |
| CCL20 | 6364 | 22.14383216 | 1.617211937 | 0.44564693 | -3.6289085 | 0.00028462 | 0.003506256 |
| KLHL35 | 283212 | 21.15817844 | 1.611841494 | 0.42931226 | -3.7544735 | 0.00017371 | 0.002397209 |
| RHCG | 51458 | 21.04340477 | 1.590905387 | 0.44680847 | -3.5605981 | 0.00037001 | 0.004279407 |
| TEX48 | 100505478 | 25.16979784 | 1.573470802 | 0.39039865 | -4.0304207 | 5.57E-05 | 0.000966652 |
| CLDN7 | 1366 | 408.7767271 | 1.552425951 | 0.12645004 | -12.276991 | 1.20E-34 | 6.05E-31 |
| BAIAP2L2 | 80115 | 26.36411284 | 1.542749432 | 0.42765982 | -3.607422 | 0.00030925 | 0.003731357 |
| CALHM5 | 254228 | 19.76155454 | 1.489224697 | 0.45235083 | -3.2921896 | 0.00099411 | 0.009541931 |
| ACP7 | 390928 | 76.35662391 | 1.481918959 | 0.23754092 | -6.238584 | 4.42E-10 | 4.57E-08 |
| LINC01116 | 375295 | 30.82930352 | 1.466103627 | 0.35944621 | -4.0787845 | 4.53E-05 | 0.000820007 |
| MESP2 | 145873 | 20.07124093 | 1.461196886 | 0.44016472 | -3.3196593 | 0.00090127 | 0.008865664 |
| HES2 | 54626 | 36.46603671 | 1.448925255 | 0.34423062 | -4.2091702 | 2.56E-05 | 0.000527677 |
| INSYN1 | 388135 | 42.31556738 | 1.424989157 | 0.31758228 | -4.486992 | 7.22E-06 | 0.000184821 |
| KLRF1 | 51348 | 27.12628109 | 1.389216249 | 0.39839069 | -3.48707 | 0.00048834 | 0.0053136 |
| LIPH | 200879 | 279.439512 | 1.38494443 | 0.13569461 | -10.206333 | 1.86E-24 | 3.50E-21 |
| GJA1 | 2697 | 25.91307546 | 1.384095067 | 0.38155289 | -3.6275314 | 0.00028614 | 0.003520155 |
| TMC4 | 147798 | 83.19146686 | 1.37507354 | 0.27343464 | -5.028893 | 4.93E-07 | 2.04E-05 |
| ENO4 | 387712 | 25.19348472 | 1.374523012 | 0.39833239 | -3.4506935 | 0.00055915 | 0.005946621 |
| FRMD4B | 23150 | 42.1681164 | 1.370055634 | 0.3093908 | -4.4282365 | 9.50E-06 | 0.000230556 |
| IL1RL1 | 9173 | 55.05120475 | 1.363960711 | 0.28844589 | -4.7286536 | 2.26E-06 | 7.10E-05 |
| SERPINB5 | 5268 | 87.28331549 | 1.363911806 | 0.21867166 | -6.2372592 | 4.45E-10 | 4.57E-08 |
| SYNE4 | 163183 | 60.56675234 | 1.356310187 | 0.2883243 | -4.7041134 | 2.55E-06 | 7.79E-05 |
| BCL2A1 | 597 | 36.17280179 | 1.350262575 | 0.32525266 | -4.1514267 | 3.30E-05 | 0.000644868 |
| LAD1 | 3898 | 56.56492221 | 1.338251326 | 0.2730649 | -4.9008543 | 9.54E-07 | 3.52E-05 |
| SMIM24 | 284422 | 24.80006715 | 1.336496473 | 0.385496 | -3.4669529 | 0.00052639 | 0.005630064 |
| FGFBP1 | 9982 | 258.8525854 | 1.32832119 | 0.14789658 | -8.9814193 | 2.67E-19 | 2.69E-16 |
| KCNQ2 | 3785 | 47.11011029 | 1.325689709 | 0.28364837 | -4.6737081 | 2.96E-06 | 8.91E-05 |
| ADGRE2 | 30817 | 53.07536437 | 1.32539148 | 0.27993183 | -4.7346937 | 2.19E-06 | 6.90E-05 |
| C1QTNF1 | 114897 | 30.54785839 | 1.316786382 | 0.37840125 | -3.479868 | 0.00050166 | 0.005427691 |
| SH3TC2 | 79628 | 77.23430256 | 1.31462071 | 0.23123706 | -5.6851643 | 1.31E-08 | 8.83E-07 |

| | | | | | | | |
|---|---|---|---|---|---|---|---|
| PTGER1 | 5731 | 87.3473274 | 1.30450354 | 0.22316773 | -5.8453951 | 5.05E-09 | 3.72E-07 |
| SUSD3 | 203328 | 53.45991928 | 1.303690798 | 0.29104026 | -4.4794175 | 7.48E-06 | 0.00018989 |
| EPS8L1 | 54869 | 39.09838468 | 1.298901279 | 0.31366253 | -4.1410789 | 3.46E-05 | 0.000666156 |
| RGS11 | 8786 | 99.38296809 | 1.296040131 | 0.22719658 | -5.7044878 | 1.17E-08 | 7.96E-07 |
| SYT16 | 83851 | 24.37139607 | 1.291526852 | 0.39287204 | -3.2873982 | 0.00101118 | 0.009662931 |
| ADAMTS10 | 81794 | 66.79513446 | 1.279209405 | 0.26927414 | -4.7505839 | 2.03E-06 | 6.48E-05 |
| FSTL1 | 11167 | 273.6856945 | 1.270633268 | 0.13209215 | -9.6192942 | 6.63E-22 | 8.01E-19 |
| GNG2 | 54331 | 36.70623625 | 1.263249169 | 0.32212215 | -3.9216464 | 8.79E-05 | 0.00139217 |
| PRTN3 | 5657 | 26.73468759 | 1.261026304 | 0.38112933 | -3.3086572 | 0.00093745 | 0.009132064 |
| INSC | 387755 | 26.12036959 | 1.251816994 | 0.37855586 | -3.3068224 | 0.00094361 | 0.009162474 |
| PRRG2 | 5639 | 52.0153423 | 1.24521457 | 0.27735858 | -4.4895476 | 7.14E-06 | 0.000183552 |
| TMEM92 | 162461 | 51.47978062 | 1.24199206 | 0.29807346 | -4.1667314 | 3.09E-05 | 0.000610302 |
| AATK | 9625 | 105.9125476 | 1.229507595 | 0.20855265 | -5.8954302 | 3.74E-09 | 2.92E-07 |
| DNER | 92737 | 118.711709 | 1.229463263 | 0.18907993 | -6.5023468 | 7.91E-11 | 1.01E-08 |
| QPCT | 25797 | 33.37953613 | 1.227254865 | 0.33768865 | -3.6342793 | 0.00027876 | 0.003454683 |
| CARMIL3 | 90668 | 38.07731103 | 1.213134083 | 0.31868143 | -3.8067297 | 0.00014082 | 0.002029834 |
| TERT | 7015 | 79.54932962 | 1.197803605 | 0.2279637 | -5.2543612 | 1.49E-07 | 7.36E-06 |
| ADM | 133 | 230.2165564 | 1.187228883 | 0.14710669 | -8.0705295 | 7.00E-16 | 3.20E-13 |
| TMEM61 | 199964 | 70.1777136 | 1.186339047 | 0.24539596 | -4.8343869 | 1.34E-06 | 4.63E-05 |
| FAM83E | 54854 | 47.20990941 | 1.185858525 | 0.29022394 | -4.0860121 | 4.39E-05 | 0.000801634 |
| FOXL1 | 2300 | 44.74835767 | 1.182475765 | 0.29450783 | -4.0150911 | 5.94E-05 | 0.001021097 |
| AZU1 | 566 | 52.1387685 | 1.181461361 | 0.30766937 | -3.8400357 | 0.00012302 | 0.001819429 |
| PKP1 | 5317 | 33.69265487 | 1.180476485 | 0.33251659 | -3.5501281 | 0.00038504 | 0.004409278 |
| MAPK15 | 225689 | 44.48308856 | 1.178291887 | 0.29682631 | -3.9696342 | 7.20E-05 | 0.001193383 |
| ABCG1 | 9619 | 75.99991908 | 1.174243707 | 0.23249065 | -5.0507136 | 4.40E-07 | 1.85E-05 |
| DAW1 | 164781 | 704.0963443 | 1.17040429 | 0.10050349 | -11.645409 | 2.42E-31 | 7.30E-28 |
| MESP1 | 55897 | 125.9850592 | 1.168758284 | 0.19760234 | -5.9146986 | 3.32E-09 | 2.65E-07 |
| SLC19A3 | 80704 | 160.5869051 | 1.168087804 | 0.1629161 | -7.1698734 | 7.51E-13 | 1.66E-10 |
| PLEKHG6 | 55200 | 89.9020547 | 1.165825029 | 0.21115683 | -5.5211335 | 3.37E-08 | 2.10E-06 |
| F12 | 2161 | 246.6795035 | 1.159745718 | 0.14480005 | -8.009291 | 1.15E-15 | 4.97E-13 |
| KIAA1549L | 25758 | 78.35250561 | 1.157360534 | 0.23754431 | -4.8721879 | 1.10E-06 | 3.95E-05 |
| NPW | 283869 | 105.8936479 | 1.154700486 | 0.22086826 | -5.2280054 | 1.71E-07 | 8.30E-06 |
| PRPH | 5630 | 60.60847336 | 1.150684085 | 0.25847786 | -4.4517705 | 8.52E-06 | 0.000210745 |
| WAKMAR2 | 100130476 | 71.63023732 | 1.14251858 | 0.22859593 | -4.997983 | 5.79E-07 | 2.30E-05 |
| TGFBR3L | 100507588 | 148.5738217 | 1.139567267 | 0.17618308 | -6.4680859 | 9.93E-11 | 1.23E-08 |
| MLPH | 79083 | 266.6372247 | 1.137833538 | 0.14459003 | -7.8693773 | 3.56E-15 | 1.25E-12 |
| MYPN | 84665 | 242.4723645 | 1.136844128 | 0.15528838 | -7.3208579 | 2.46E-13 | 6.17E-11 |
| DENND1C | 79958 | 64.11457106 | 1.131167045 | 0.26369848 | -4.289623 | 1.79E-05 | 0.000386812 |
| ANGPTL4 | 51129 | 516.9217151 | 1.130554621 | 0.11758255 | -9.6149863 | 6.91E-22 | 8.01E-19 |
| SPRY4 | 81848 | 73.5328524 | 1.112077335 | 0.26379917 | -4.2156211 | 2.49E-05 | 0.000514221 |
| TMEM121 | 80757 | 133.3329789 | 1.1005935 | 0.1851874 | -5.9431337 | 2.80E-09 | 2.29E-07 |
| KRT86 | 3892 | 191.2190306 | 1.097548034 | 0.16769116 | -6.5450559 | 5.95E-11 | 8.07E-09 |
| PALM3 | 342979 | 74.33614676 | 1.096101271 | 0.22704921 | -4.8275935 | 1.38E-06 | 4.73E-05 |
| PTH1R | 5745 | 149.7158525 | 1.093244644 | 0.16870737 | -6.4801239 | 9.16E-11 | 1.15E-08 |
| WNT10B | 7480 | 81.82237985 | 1.0914269 | 0.21813086 | -5.003542 | 5.63E-07 | 2.25E-05 |
| CCDC74B | 91409 | 98.15432006 | 1.090156528 | 0.22300379 | -4.8885113 | 1.02E-06 | 3.72E-05 |
| PSG1 | 5669 | 41.08828565 | 1.086438461 | 0.31590135 | -3.4391701 | 0.0005835 | 0.006148779 |
| CES3 | 23491 | 105.524497 | 1.084469211 | 0.20160109 | -5.3792824 | 7.48E-08 | 4.10E-06 |
| HAAO | 23498 | 52.58847493 | 1.068841739 | 0.27269592 | -3.919537 | 8.87E-05 | 0.001401154 |

| | | | | | | | |
|---|---|---|---|---|---|---|---|
| STAP2 | 55620 | 59.24706745 | 1.057775929 | 0.25205649 | -4.1965827 | 2.71E-05 | 0.000548128 |
| RASGRP2 | 10235 | 58.4144214 | 1.054615843 | 0.25257484 | -4.1754588 | 2.97E-05 | 0.000592022 |
| RASD2 | 23551 | 104.2740408 | 1.052837372 | 0.19916553 | -5.2862429 | 1.25E-07 | 6.42E-06 |
| STC1 | 6781 | 715.4362869 | 1.051846598 | 0.09871355 | -10.655544 | 1.64E-26 | 3.54E-23 |
| COL9A3 | 1299 | 86.02519414 | 1.050265829 | 0.21766091 | -4.8252386 | 1.40E-06 | 4.77E-05 |
| KCTD19 | 146212 | 133.2238846 | 1.049612917 | 0.1795657 | -5.8452861 | 5.06E-09 | 3.72E-07 |
| COL13A1 | 1305 | 333.3462086 | 1.048441262 | 0.13142287 | -7.9776164 | 1.49E-15 | 5.83E-13 |
| B4GALNT2 | 124872 | 75.82651354 | 1.041848865 | 0.22527115 | -4.6248658 | 3.75E-06 | 0.000106784 |
| SULT2B1 | 6820 | 62.90469693 | 1.039053937 | 0.26376156 | -3.9393683 | 8.17E-05 | 0.001315348 |
| LOXL1 | 4016 | 59.04851642 | 1.031827614 | 0.25037855 | -4.1210703 | 3.77E-05 | 0.000713067 |
| DUSP4 | 1846 | 337.8021977 | 1.029430073 | 0.12347994 | -8.3368204 | 7.63E-17 | 4.42E-14 |
| DUOX1 | 53905 | 55.20277185 | 1.028411103 | 0.2631906 | -3.9074766 | 9.33E-05 | 0.001453469 |
| RNF223 | 401934 | 61.6766655 | 1.028127207 | 0.2508176 | -4.0991031 | 4.15E-05 | 0.000765036 |
| L1CAM | 3897 | 147.1292354 | 1.026349808 | 0.17047105 | -6.0206691 | 1.74E-09 | 1.49E-07 |
| RASAL1 | 8437 | 96.76487197 | 1.025525268 | 0.2123231 | -4.830022 | 1.37E-06 | 4.70E-05 |
| TSPAN8 | 7103 | 74.67486716 | 1.013403596 | 0.25029498 | -4.048837 | 5.15E-05 | 0.000906185 |
| APLP1 | 333 | 895.159106 | 1.011559277 | 0.08952731 | -11.298891 | 1.33E-29 | 3.34E-26 |
| CCN3 | 4856 | 51.32140079 | 1.001808508 | 0.28125864 | -3.5618764 | 0.00036821 | 0.004265166 |
| DNAJC12 | 56521 | 68.02304609 | 0.998112957 | 0.26787451 | -3.7260468 | 0.00019451 | 0.002614817 |
| FBXO6 | 26270 | 141.6609501 | 0.996914984 | 0.19183057 | -5.1968515 | 2.03E-07 | 9.61E-06 |
| TMEM74B | 55321 | 83.7987934 | 0.996606645 | 0.22047436 | -4.5202837 | 6.18E-06 | 0.000164721 |
| WIPF3 | 644150 | 230.5592519 | 0.992347567 | 0.14056024 | -7.0599449 | 1.67E-12 | 3.35E-10 |
| MDFI | 4188 | 407.501634 | 0.987207414 | 0.12376914 | -7.9761999 | 1.51E-15 | 5.83E-13 |
| AOX1 | 316 | 267.990915 | 0.986157108 | 0.1433555 | -6.879102 | 6.02E-12 | 1.01E-09 |
| ANXA9 | 8416 | 64.46002649 | 0.984659939 | 0.23776464 | -4.1413221 | 3.45E-05 | 0.000666156 |
| PIP5KL1 | 138429 | 795.0865471 | 0.984086474 | 0.26612169 | -3.6978816 | 0.00021741 | 0.002844021 |
| HYI | 81888 | 1004.496251 | 0.983378249 | 0.10019549 | -9.8145957 | 9.74E-23 | 1.63E-19 |
| TMEM151A | 256472 | 63.75800639 | 0.982006016 | 0.25093574 | -3.9133764 | 9.10E-05 | 0.001425768 |
| G0S2 | 50486 | 476.1092333 | 0.980988167 | 0.13843367 | -7.0863407 | 1.38E-12 | 2.92E-10 |
| CITED4 | 163732 | 45.95955322 | 0.980709831 | 0.28725559 | -3.414067 | 0.00064001 | 0.006656263 |
| TSPAN18 | 90139 | 230.3174438 | 0.977636407 | 0.14157438 | -6.9054615 | 5.00E-12 | 8.77E-10 |
| NKD2 | 85409 | 907.3636535 | 0.97174049 | 0.11359917 | -8.5541163 | 1.19E-17 | 8.14E-15 |
| SCARA5 | 286133 | 168.2033401 | 0.9707754 | 0.16664693 | -5.8253423 | 5.70E-09 | 4.15E-07 |
| KCNN4 | 3783 | 441.8956473 | 0.970399152 | 0.11981812 | -8.0989349 | 5.54E-16 | 2.61E-13 |
| LGALS8-AS1 | 100287902 | 63.75948556 | 0.964251441 | 0.24407532 | -3.9506307 | 7.79E-05 | 0.001271254 |
| SSPN | 8082 | 46.88251539 | 0.96058585 | 0.28368406 | -3.3861114 | 0.00070891 | 0.007257617 |
| CHST13 | 166012 | 105.8485718 | 0.955003518 | 0.21270506 | -4.4898016 | 7.13E-06 | 0.000183552 |
| HES7 | 84667 | 361.1279759 | 0.953185027 | 0.1331864 | -7.1567747 | 8.26E-13 | 1.80E-10 |
| NEFL | 4747 | 200.7455509 | 0.953014683 | 0.14642085 | -6.5087364 | 7.58E-11 | 9.93E-09 |
| ACE | 1636 | 87.38067234 | 0.9503404 | 0.21568934 | -4.406061 | 1.05E-05 | 0.000251407 |
| ARTN | 9048 | 74.16282496 | 0.945514505 | 0.22997746 | -4.1113356 | 3.93E-05 | 0.000737337 |
| LAT2 | 7462 | 195.4935926 | 0.944088696 | 0.15347302 | -6.1514963 | 7.68E-10 | 7.56E-08 |
| CAMK2N2 | 94032 | 81.8012444 | 0.94112401 | 0.22606705 | -4.1630304 | 3.14E-05 | 0.000618661 |
| ZNF296 | 162979 | 44.1168505 | 0.939442905 | 0.28454441 | -3.3015687 | 0.00096146 | 0.009287931 |
| UNC13D | 201294 | 230.2962252 | 0.937010502 | 0.14403224 | -6.505561 | 7.74E-11 | 1.01E-08 |
| LINC02381 | 400043 | 54.97634785 | 0.935778332 | 0.25902702 | -3.6126669 | 0.00030306 | 0.003671361 |
| CA12 | 771 | 169.6710151 | 0.930606053 | 0.15759902 | -5.9048975 | 3.53E-09 | 2.78E-07 |
| CREG2 | 200407 | 64.41168355 | 0.929879574 | 0.23881498 | -3.8937239 | 9.87E-05 | 0.001519576 |
| PLEKHA6 | 22874 | 152.7473984 | 0.929368766 | 0.17681652 | -5.2561196 | 1.47E-07 | 7.32E-06 |

| | | | | | | | |
|---|---|---|---|---|---|---|---|
| CELF5 | 60680 | 77.10973701 | 0.924779148 | 0.22425403 | -4.1238018 | 3.73E-05 | 0.000708213 |
| GALNT6 | 11226 | 106.6789657 | 0.924467922 | 0.19847072 | -4.6579561 | 3.19E-06 | 9.38E-05 |
| MIR210HG | 100506211 | 305.2809665 | 0.918654646 | 0.1306354 | -7.0322032 | 2.03E-12 | 3.98E-10 |
| BSPRY | 54836 | 227.2447292 | 0.917159098 | 0.13770544 | -6.6602969 | 2.73E-11 | 3.81E-09 |
| HLA-DRB1 | 3123 | 262.5134278 | 0.916561311 | 0.15019547 | -6.1024565 | 1.04E-09 | 9.66E-08 |
| TMEM229B | 161145 | 74.70278513 | 0.91530222 | 0.2299158 | -3.9810323 | 6.86E-05 | 0.00114513 |
| TNFSF10 | 8743 | 124.1158188 | 0.914374165 | 0.1814377 | -5.0396042 | 4.66E-07 | 1.96E-05 |
| C6orf132 | 647024 | 146.2977545 | 0.911115912 | 0.16643076 | -5.4744442 | 4.39E-08 | 2.59E-06 |
| NDRG1 | 10397 | 5511.921069 | 0.91059877 | 0.09463563 | -9.6221553 | 6.45E-22 | 8.01E-19 |
| ETV4 | 2118 | 382.2606543 | 0.910135915 | 0.122825 | -7.4100217 | 1.26E-13 | 3.28E-11 |
| STC2 | 8614 | 2312.761748 | 0.90933215 | 0.0936837 | -9.7064071 | 2.83E-22 | 4.27E-19 |
| ARHGAP4 | 393 | 132.8908935 | 0.903255714 | 0.19232874 | -4.6964157 | 2.65E-06 | 8.06E-05 |
| BMPER | 168667 | 130.8648173 | 0.90209066 | 0.19304472 | -4.6729622 | 2.97E-06 | 8.91E-05 |
| CXCL8 | 3576 | 619.2939693 | 0.897782059 | 0.10487006 | -8.5608996 | 1.12E-17 | 8.04E-15 |
| EFEMP2 | 30008 | 176.5913295 | 0.895842237 | 0.17583343 | -5.0948345 | 3.49E-07 | 1.51E-05 |
| ARHGAP40 | 343578 | 93.09264182 | 0.894190187 | 0.2243495 | -3.9857018 | 6.73E-05 | 0.001126583 |
| ESRP1 | 54845 | 162.1343505 | 0.893779676 | 0.16936976 | -5.2770914 | 1.31E-07 | 6.70E-06 |
| ERFE | 151176 | 92.09975946 | 0.892687051 | 0.22960862 | -3.8878639 | 0.00010113 | 0.001545673 |
| WNT5B | 81029 | 160.0304325 | 0.887757817 | 0.16207502 | -5.47745 | 4.31E-08 | 2.56E-06 |
| PPP1R3G | 648791 | 235.5291983 | 0.882562288 | 0.15178607 | -5.8145145 | 6.08E-09 | 4.36E-07 |
| NALT1 | 101928483 | 73.82571681 | 0.882333458 | 0.22415836 | -3.9362059 | 8.28E-05 | 0.001329951 |
| TNNT1 | 7138 | 1079.679098 | 0.882065221 | 0.11185542 | -7.885762 | 3.13E-15 | 1.12E-12 |
| NLRP1 | 22861 | 68.81433374 | 0.877828642 | 0.23141733 | -3.793271 | 0.00014868 | 0.00212455 |
| IRF5 | 3663 | 300.4880596 | 0.874219343 | 0.13124575 | -6.6609346 | 2.72E-11 | 3.81E-09 |
| NGEF | 25791 | 134.4466052 | 0.871866556 | 0.17899821 | -4.8708116 | 1.11E-06 | 3.96E-05 |
| WFDC2 | 10406 | 229.5906195 | 0.866056206 | 0.16703136 | -5.1849918 | 2.16E-07 | 1.01E-05 |
| LOC101929705 | 101929705 | 97.4656173 | 0.865835587 | 0.22188554 | -3.9021722 | 9.53E-05 | 0.001479581 |
| LTBP4 | 8425 | 3622.859577 | 0.865384675 | 0.09706596 | -8.9154289 | 4.86E-19 | 4.31E-16 |
| TNFRSF14 | 8764 | 82.94775855 | 0.860947661 | 0.24328473 | -3.5388479 | 0.00040188 | 0.004563728 |
| FER1L4 | 80307 | 791.2926466 | 0.86051595 | 0.26195762 | -3.2849434 | 0.00102003 | 0.009722853 |
| MMP1 | 4312 | 124.2437872 | 0.856306634 | 0.19526006 | -4.3854675 | 1.16E-05 | 0.000271675 |
| NPAS1 | 4861 | 152.5696309 | 0.856014151 | 0.17329785 | -4.9395544 | 7.83E-07 | 2.99E-05 |
| PADI2 | 11240 | 192.5521913 | 0.851369666 | 0.14737706 | -5.7768127 | 7.61E-09 | 5.34E-07 |
| KIRREL2 | 84063 | 106.022171 | 0.847762833 | 0.21080115 | -4.0216233 | 5.78E-05 | 0.000996593 |
| LRRC34 | 151827 | 266.3317996 | 0.84771187 | 0.13317215 | -6.365534 | 1.95E-10 | 2.14E-08 |
| HK2 | 3099 | 721.8788965 | 0.841619913 | 0.09863449 | -8.5327146 | 1.43E-17 | 9.37E-15 |
| CARD11 | 84433 | 99.87209684 | 0.84009749 | 0.19674858 | -4.2699037 | 1.96E-05 | 0.00041625 |
| LYPD3 | 27076 | 750.5661199 | 0.839210796 | 0.10153214 | -8.2654693 | 1.39E-16 | 7.27E-14 |
| GGT1 | 2678 | 170.2338261 | 0.838593829 | 0.17985591 | -4.662587 | 3.12E-06 | 9.26E-05 |
| GNA15 | 2769 | 124.7685583 | 0.838135526 | 0.18643827 | -4.4955123 | 6.94E-06 | 0.000179092 |
| CEMIP | 57214 | 2550.564939 | 0.837849162 | 0.09680853 | -8.654704 | 4.94E-18 | 3.72E-15 |
| MFSD2A | 84879 | 85.05677016 | 0.832445443 | 0.22837874 | -3.6450216 | 0.00026737 | 0.003354724 |
| EML1 | 2009 | 261.5867901 | 0.831691515 | 0.1405623 | -5.9168888 | 3.28E-09 | 2.64E-07 |
| VTN | 7448 | 161.051236 | 0.831650907 | 0.17712339 | -4.6953196 | 2.66E-06 | 8.09E-05 |
| LHX2 | 9355 | 218.2030754 | 0.831502868 | 0.14037207 | -5.9235636 | 3.15E-09 | 2.55E-07 |
| NUDT8 | 254552 | 154.9426855 | 0.829421617 | 0.16594117 | -4.998287 | 5.78E-07 | 2.30E-05 |
| MAPK8IP2 | 23542 | 589.6770752 | 0.828204191 | 0.11906053 | -6.9561606 | 3.50E-12 | 6.35E-10 |
| GRIN1 | 2902 | 92.24683377 | 0.827697065 | 0.21354714 | -3.8759455 | 0.00010621 | 0.00160865 |
| SPDEF | 25803 | 79.69208905 | 0.827523581 | 0.23811556 | -3.4753024 | 0.00051028 | 0.005484941 |

| | | | | | | | |
|---|---|---|---|---|---|---|---|
| MACROD1 | 28992 | 302.8167764 | 0.823241513 | 0.16012796 | -5.141148 | 2.73E-07 | 1.23E-05 |
| C19orf81 | 342918 | 75.7654431 | 0.821408689 | 0.22330828 | -3.678362 | 0.00023474 | 0.003024431 |
| CARD6 | 84674 | 120.7787593 | 0.816560102 | 0.18727216 | -4.3602855 | 1.30E-05 | 0.000297257 |
| EVI2B | 2124 | 86.27110041 | 0.812457486 | 0.21259582 | -3.8216061 | 0.00013259 | 0.001926771 |
| KISS1R | 84634 | 81.61187624 | 0.812337315 | 0.22750783 | -3.5705906 | 0.00035618 | 0.004161145 |
| IGDCC4 | 57722 | 387.5145404 | 0.810478647 | 0.12033341 | -6.7352753 | 1.64E-11 | 2.44E-09 |
| DNAAF3 | 352909 | 124.4113965 | 0.809810314 | 0.18669963 | -4.3375036 | 1.44E-05 | 0.000324625 |
| LOXL4 | 84171 | 292.7765977 | 0.8094291 | 0.12631602 | -6.4079686 | 1.47E-10 | 1.70E-08 |
| PAQR5 | 54852 | 130.8065966 | 0.807257329 | 0.17634129 | -4.5778122 | 4.70E-06 | 0.000129686 |
| HSPB8 | 26353 | 112.9524042 | 0.807071955 | 0.19314033 | -4.1786816 | 2.93E-05 | 0.000587577 |
| NRARP | 441478 | 152.3778337 | 0.80559869 | 0.16610383 | -4.8499705 | 1.23E-06 | 4.31E-05 |
| SPTBN4 | 57731 | 112.7296362 | 0.803707662 | 0.19571364 | -4.106549 | 4.02E-05 | 0.000745787 |
| TLL2 | 7093 | 74.61128171 | 0.802517438 | 0.24424284 | -3.2857357 | 0.00101716 | 0.009703555 |
| ADAMTS7 | 11173 | 254.4261637 | 0.801254598 | 0.14863247 | -5.3908448 | 7.01E-08 | 3.90E-06 |
| FAM131C | 348487 | 286.1934338 | 0.797796753 | 0.13046101 | -6.115212 | 9.64E-10 | 9.14E-08 |
| CACNG8 | 59283 | 122.0639063 | 0.793757395 | 0.18997696 | -4.1781772 | 2.94E-05 | 0.0005881 |
| PPFIA3 | 8541 | 239.2503444 | 0.793684794 | 0.16307982 | -4.8668487 | 1.13E-06 | 4.02E-05 |
| NEIL1 | 79661 | 246.8892769 | 0.786665229 | 0.15634154 | -5.0317096 | 4.86E-07 | 2.02E-05 |
| PGF | 5228 | 100.2225987 | 0.786223869 | 0.22314421 | -3.523389 | 0.00042607 | 0.00476321 |
| SECTM1 | 6398 | 500.3181305 | 0.785272862 | 0.11172778 | -7.0284475 | 2.09E-12 | 4.03E-10 |
| STK31 | 56164 | 304.5528069 | 0.783875451 | 0.13342086 | -5.875209 | 4.22E-09 | 3.23E-07 |
| CRIP1 | 1396 | 3658.828319 | 0.783628413 | 0.10106702 | -7.7535525 | 8.94E-15 | 2.92E-12 |
| ST8SIA6 | 338596 | 71.02272435 | 0.783245793 | 0.23388035 | -3.3489167 | 0.00081128 | 0.008139826 |
| KRT15 | 3866 | 71.02812729 | 0.779300657 | 0.23574865 | -3.3056421 | 0.00094759 | 0.009195227 |
| CXCL5 | 6374 | 192.9660684 | 0.779012539 | 0.15693046 | -4.9640619 | 6.90E-07 | 2.66E-05 |
| APOE | 348 | 609.7918727 | 0.778919055 | 0.1124174 | -6.9288121 | 4.24E-12 | 7.61E-10 |
| LINC00957 | 255031 | 153.322615 | 0.774428044 | 0.17971361 | -4.3092342 | 1.64E-05 | 0.000360933 |
| INAVA | 55765 | 150.2835474 | 0.768813713 | 0.17032476 | -4.5138107 | 6.37E-06 | 0.000168343 |
| RCOR2 | 283248 | 86.42220791 | 0.767655419 | 0.21809518 | -3.5198183 | 0.00043184 | 0.004809953 |
| COL16A1 | 1307 | 74.73285051 | 0.766268744 | 0.22285204 | -3.4384641 | 0.00058502 | 0.006156644 |
| NID2 | 22795 | 206.6733637 | 0.76461715 | 0.14281526 | -5.3538897 | 8.61E-08 | 4.63E-06 |
| NFASC | 23114 | 123.8104847 | 0.763613986 | 0.18356468 | -4.1599178 | 3.18E-05 | 0.000624703 |
| NPTX1 | 4884 | 107.4769931 | 0.760881802 | 0.21111349 | -3.6041363 | 0.00031319 | 0.003769819 |
| BDH1 | 622 | 269.3464489 | 0.760550976 | 0.12970928 | -5.8635046 | 4.53E-09 | 3.41E-07 |
| HOXC5 | 3222 | 126.4353462 | 0.760474207 | 0.18689236 | -4.0690492 | 4.72E-05 | 0.000848908 |
| ALDH1L2 | 160428 | 355.8860086 | 0.757370394 | 0.12913045 | -5.8651572 | 4.49E-09 | 3.40E-07 |
| NKX2-8 | 26257 | 241.1288378 | 0.756972873 | 0.1630436 | -4.6427636 | 3.44E-06 | 9.91E-05 |
| SLC7A5 | 8140 | 9021.500531 | 0.756704409 | 0.08474358 | -8.9293425 | 4.29E-19 | 4.04E-16 |
| SLC37A1 | 54020 | 294.736694 | 0.756059665 | 0.12666779 | -5.968839 | 2.39E-09 | 2.00E-07 |
| GRIN3B | 116444 | 196.2720497 | 0.755211105 | 0.16037971 | -4.7088943 | 2.49E-06 | 7.69E-05 |
| NES | 10763 | 420.3418837 | 0.754343306 | 0.12689539 | -5.9446076 | 2.77E-09 | 2.28E-07 |
| NAGS | 162417 | 517.4634759 | 0.751533754 | 0.11512603 | -6.5279223 | 6.67E-11 | 8.89E-09 |
| SELENOP | 6414 | 177.8911917 | 0.751386044 | 0.1802385 | -4.1688433 | 3.06E-05 | 0.000605469 |
| CORO6 | 84940 | 318.5474265 | 0.751367427 | 0.1265908 | -5.935403 | 2.93E-09 | 2.39E-07 |
| SH2B2 | 10603 | 118.947637 | 0.750363275 | 0.18284776 | -4.1037597 | 4.06E-05 | 0.000751632 |
| PPP1R16B | 26051 | 138.950063 | 0.749104055 | 0.18163824 | -4.1241539 | 3.72E-05 | 0.000708024 |
| TNNI3 | 7137 | 132.2212764 | 0.743780309 | 0.17812449 | -4.1756207 | 2.97E-05 | 0.000592022 |
| SPINT1 | 6692 | 1427.904482 | 0.739873537 | 0.09321818 | -7.9370087 | 2.07E-15 | 7.80E-13 |
| TMEM30B | 161291 | 208.6124251 | 0.739506893 | 0.15533709 | -4.760659 | 1.93E-06 | 6.23E-05 |

| | | | | | | | |
|---|---|---|---|---|---|---|---|
| HOXB9 | 3219 | 719.7567467 | 0.733324266 | 0.09371731 | -7.8248538 | 5.08E-15 | 1.73E-12 |
| DGCR5 | 26220 | 415.2265283 | 0.733311477 | 0.1171613 | -6.2589906 | 3.87E-10 | 4.08E-08 |
| CYS1 | 192668 | 124.9665949 | 0.729862903 | 0.19739312 | -3.6975094 | 0.00021773 | 0.002845723 |
| MSI1 | 4440 | 174.0078681 | 0.728301602 | 0.1549027 | -4.7016714 | 2.58E-06 | 7.87E-05 |
| NOXA1 | 10811 | 438.5813501 | 0.726074836 | 0.13697035 | -5.3009637 | 1.15E-07 | 6.01E-06 |
| STEAP1B | 256227 | 85.33303595 | 0.722806195 | 0.21503853 | -3.3612869 | 0.0007758 | 0.007851805 |
| CSGALNACT1 | 55790 | 492.2332612 | 0.719983981 | 0.10552991 | -6.8225584 | 8.94E-12 | 1.39E-09 |
| HHIPL1 | 84439 | 166.094212 | 0.719110143 | 0.16820727 | -4.2751431 | 1.91E-05 | 0.000408891 |
| LGI3 | 203190 | 133.9902869 | 0.714822984 | 0.18299302 | -3.9062856 | 9.37E-05 | 0.001459139 |
| EPOP | 100170841 | 515.2525388 | 0.713143084 | 0.11825305 | -6.0306529 | 1.63E-09 | 1.41E-07 |
| PRR29 | 92340 | 113.1888916 | 0.707842237 | 0.20369721 | -3.4749727 | 0.00051091 | 0.005487772 |
| CPT1C | 126129 | 169.2052761 | 0.707301359 | 0.17491735 | -4.0436318 | 5.26E-05 | 0.00092117 |
| MAP3K7CL | 56911 | 173.7332349 | 0.707210153 | 0.15334903 | -4.611768 | 3.99E-06 | 0.000112886 |
| DDIT4 | 54541 | 4612.761209 | 0.706918936 | 0.09167448 | -7.7111858 | 1.25E-14 | 3.91E-12 |
| FBXL16 | 146330 | 274.4928042 | 0.7055123 | 0.14992743 | -4.7056918 | 2.53E-06 | 7.75E-05 |
| EGFL7 | 51162 | 1431.442486 | 0.704617691 | 0.09383432 | -7.5091682 | 5.95E-14 | 1.79E-11 |
| HAPLN3 | 145864 | 121.7735612 | 0.704242028 | 0.19496599 | -3.6121276 | 0.0003037 | 0.003674802 |
| PTPRU | 10076 | 298.2059944 | 0.704162355 | 0.13998469 | -5.0302812 | 4.90E-07 | 2.03E-05 |
| PTK6 | 5753 | 280.9850334 | 0.700013536 | 0.13569735 | -5.1586383 | 2.49E-07 | 1.14E-05 |
| LY6K | 54742 | 3321.602029 | 0.699376851 | 0.09726835 | -7.1901787 | 6.47E-13 | 1.46E-10 |
| LINC01588 | 283551 | 302.3285816 | 0.694576518 | 0.13260986 | -5.2377442 | 1.63E-07 | 7.90E-06 |
| GAMT | 2593 | 962.5101188 | 0.690666562 | 0.11701248 | -5.9025034 | 3.58E-09 | 2.81E-07 |
| ARHGEF37 | 389337 | 377.9249812 | 0.690610008 | 0.11263038 | -6.1316492 | 8.70E-10 | 8.46E-08 |
| ABHD1 | 84696 | 124.401849 | 0.690258138 | 0.18061774 | -3.8216519 | 0.00013256 | 0.001926771 |
